# Gray Matter Volumetric Correlates of Attention Deficit and Hyperactivity Traits in Emerging Adolescents

**DOI:** 10.1101/2022.05.16.492088

**Authors:** Clara S. Li, Yu Chen, Jaime S. Ide

**Author notes:** Correspondence: Dr. Yu Chen, Connecticut Mental Health Center S110A, 34 Park Street, New Haven, CT 06519, U.S.A.; or Dr. Jaime S. Ide, Connecticut Mental Health Center S110, 34 Park Street, New Haven, CT 06519, U.S.A.

## Abstract

Previous research has demonstrated reduction in cortical and subcortical, including basal ganglia (BG), gray matter volumes (GMV) in individuals with attention deficit hyperactivity disorder (ADHD), a neurodevelopmental condition that is more prevalent in males than in females. However, the volumetric deficits vary across studies. Whether volumetric reductions are more significant in males than females; to what extent these neural markers are heritable and relate to cognitive dysfunction in ADHD remain unclear. To address these questions, we followed published routines and performed voxel-based morphometry analysis of a data set (n = 11,502; 5,464 girls, 9-10 years) curated from the Adolescent Brain Cognition Development project, a population-based study of typically developing children. Of the sample, 634 and 2,826 were identified as monozygotic twins (MZ) and dizygotic twins/siblings (DZ), respectively. In linear regressions, a cluster in the hypothalamus showed larger GMV, and bilateral caudate and putamen, lateral orbitofrontal and occipital cortex showed smaller GMVs, in correlation with higher ADHD scores in girls and boys combined. When examined separately, boys relative to girls showed more widespread (including BG) and stronger associations between GMV deficits and ADHD scores. ADHD traits and the volumetric correlates demonstrated heritability estimates (*a^2^*) between 0.59 and 0.79, replicating prior findings of the genetic basis of ADHD. Further, ADHD traits and the volumetric correlates (except for the hypothalamus) were each negatively and positively correlated with N-back performance. Together, these findings confirm volumetric deficits in children with more prominent ADHD traits. Highly heritable in both girls and boys and potentially more significant in boys than in girls, the structural deficits underlie diminished capacity in working memory and potentially other cognitive deficits in ADHD.

## Introduction

Attention deficit hyperactivity disorder (ADHD) is one of the most common neurodevelopmental disorders, with a prevalence rate of approximately 7% in children and 4.4% in adults^1–3^. ADHD is associated with inattention, impulsivity, hyperactivity, and an array of cognitive deficits with varying degrees of severity^4, 5^. For instance, as compared to healthy controls, individuals with ADHD showed lower capacity of working memory^6–9^, a key cognitive function that is closely associated with fluid intelligence^10–12^. Extensive research has focused on identifying the neural markers of ADHD, and many studies demonstrated gray matter volume (GMV) reduction in cortical and subcortical structures, including the basal ganglia (BG), in adults and children with ADHD^13, 14^.

A number of meta-analyses aimed to investigate the consistency of the volumetric reductions in ADHD. ADHD relative to typically developing children showed global volumetric reductions, with the cerebellum, splenium of the corpus callosum, and right caudate showing the most significant differences^15^. Other studies identified GMV reduction more specifically in the BG^16, 17^, and the volumetric deficits appeared to normalize with age and treatment with stimulant medications^17^. A meta-analysis of 11 studies comprising 320 ADHD cases and 288 controls showed reduction of the right globus pallidus and putamen, as well as bilateral caudate volumes in children with ADHD, and reduction of anterior cingulate cortex (ACC) volume in adults with ADHD^18^. A higher percentage of treated relative to non-treated participants were associated with fewer deficits, suggesting a remediating effect of the medications. The ENIGMA ADHD Working Group reported smaller volumes of the accumbens, amygdala, caudate, putamen, and hippocampus in ADHD cases at 4–63 years of age, relative to controls^19^. Effect sizes were highest in children, whereas case-control differences were not present in adults. Thus, it appears that regional volume reduction is more prominent in children than in adults with ADHD and neural development and/or chronic treatment with medications may account for the differences.

On the other hand, many other studies have described smaller volumes in the frontal cortex, temporal cortex, insula, and cerebellum in ADHD children or adults^20–29^ but did not necessarily implicate the BG. To confirm BG volumetric reduction as a neural marker of ADHD would require the investigation of a large sample of children within a limited age range and the absence of the effects of prolonged medications.

Previous work has also studied sex differences in ADHD traits and GMVs in association with ADHD. ADHD is more commonly observed in boys than girls^30, 31^. Earlier studies have suggested that boys with ADHD showed lower GMVs in the ACC and BG, whereas girls with ADHD exhibited higher GMV in the ACC and no difference in the BG, as compared to typically developing children^32–34^. Therefore, it would be critical to consider sex differences in examining volumetric reduction in the BG and other cortical and subcortical structures as neural markers of ADHD.

ADHD is highly heritable^35^, with 60 to 80% of variance in ADHD risk attributed to genetic effects^36–41^. Previous work has also suggested the heritability of the volumetric markers of ADHD. Alterations in total and BG GMV have been observed not only in individuals with ADHD but also in their unaffected siblings and relatives, suggesting genetic and shared environmental risks^42, 43^. Individuals with ADHD showed reduction in total and regional cortical GMVs, and their unaffected siblings showed intermediate GMVs between individuals with ADHD and healthy controls^25, 44^. Indeed, recent studies have aimed specifically to characterize the molecular genetic bases of the volumetric features of ADHD^45–48^. Studies have reported evidence of global pleiotropy for variants affecting ADHD risk for total intracranial and subcortical regional volumes^49^. Another ENIGMA study also showed shared genetic heritability with brain structural correlates but did not report the exact heritability estimates of ADHD or of the structural correlates^50^.

On the other hand, affected twins had significantly smaller caudate than their unaffected co-twins^51^, suggesting unique environmental influences. Further, genetic and environmental risk factors may contribute to distinct regional volume losses^52^, and the sex difference in ADHD symptoms may result from the different genetic and environmental influences on brain structures^53^.

The present study aimed to examine sex-shared and specific volumetric markers of ADHD traits for the whole brain. Because of the focus of previous studies on the BG, we also performed ROI analyses specifically of the caudate, putamen, and pallidum. Using a large data set of children (9 to 10 years) of the Adolescent Brain Cognition Development (ABCD) study, we employed voxel-based morphometry to identify GMV correlates of ADHD in girls and boys combined as well as separately. We evaluated how the volumetric deficits contribute to cognitive dysfunction, as evaluated in an N-back task, and assessed sex differences in the relationships between volumetric and cognitive deficits. Finally, we estimated the heritability of the neural and behavioral markers and examined sex differences in the heritability estimates. The overall goal was to characterize structural brain alterations during an early and prodromal stage of ADHD, as with many of the ENIGMA studies of ADHD^19, 54^, and to inform longitudinal research of later releases of the ABCD data.

## Methods

### Dataset

The ABCD Release 2.0 cohort comprised 11,601 children; however, 99 subjects were not included in the current study because of questionable image quality or poor image segmentation (details in the section on *Voxel-based morphometry*). Thus, the current sample consisted of 11,502 subjects (5,464 girls, age 9 to 10 years). Of the 11,502 children, 634 and 2,826 were identified as monozygotic twins (MZ) and dizygotic twins/siblings (DZ), respectively. The ABCD data were collected from 21 research sites across the country. Children participated at baseline and follow-up assessments over a period of 10 years. All recruitment procedures and informed consent forms, including consent to share de-identified data, were approved by the Institutional Review Board of the University of California San Diego with study number 160091. Informed consent from parent and assent from child under 18 years old was obtained from all individuals prior to participation. The consortium workgroups established standardized assessments of physical and mental health, neurocognition, substance use, culture, and environment, as well as multimodal structural and functional brain imaging and bioassay protocols (https://abcdstudy.org/). Structural magnetic resonance imaging (MRI) data were acquired using an optimized protocol for 3T machines, including Siemens Prisma, GE 750 and Philips, with voxel size 1 mm isotropic^55^. We have obtained permission from the ABCD to use the Open and Restricted Access data for the current study. All methods were performed in accordance with ABCD Bioethics and Medical Oversight Guidelines and Procedures.

### Assessments and cognitive test

The children were assessed with the ABCD Parent Child Behavior Checklist (CBCL) from the Achenbach System of Empirically Based Assessment^56^ for dimensional psychopathology and adaptive functioning. The CBCL is widely used to identify ADHD and other problem behaviors in children^57, 58^. We retrieved the “cbcl_scr_dsm5_adhd_t” values with the *t*-score in the DSM-5 scale (ADHD score, hereafter) and normed by sex, age, informant and ethnicity^59^. Seven ADHD items in the CBCL DSM-oriented scale were used in the ABCD study: #cbcl_q04_p -Fails to finish things they start; #cbcl_q08_p - Can’t concentrate, can’t pay attention for long; #cbcl_q10_p - Can’t sit still, restless, or hyperactive; #cbcl_q41_p - Impulsive or acts without thinking; #cbcl_q78_p - Inattentive or easily distracted; #cbcl_q93_p - Talks too much; and #cbcl_q104_p - Unusually loud. Each item’s response was rated on a scale 0= Not True, 1 = Somewhat or Sometimes Ture, and 2 = Very True or Often True. The sum of raw scores was transformed to normalized *T* scores (range 50-70) based on the percentiles of national normative sample of nonreferred populations^60^. Investigators have used different CBCL cutoff scores for a diagnosis of ADHD^58^. Here, for instance, with a cutoff of 65 (ADHD *T* score), 451 (7.5 %) boys and 272 (5%) girls would be considered to have a diagnosis of ADHD. With a cutoff of 70 or the 98^th^ percentile of national normative samples,193 (3.2 %) boys and 75 (1.4%) girls would be considered to have a diagnosis of ADHD.

Participants performed an N-Back task^55, 61^. Briefly, there were four blocks each with 2- and 0-back conditions in each run, with a total of 2 runs. Participants were required to respond to a set of stimuli of emotional faces or places. Each block consisted of 10 trials (2.5 s each) and 4 fixation trials (15 s each). At each trial, participants were instructed to respond to whether the picture was a “match” or “no-match” of a pre-specified target (0-back) or the stimulus shown two trials back (2-back). Here, we employed the accuracy rate of 2-back trials as a measure of the capacity of working memory.

### Customized pediatric template construction

In order to perform voxel-based morphometry with appropriate templates, we constructed customized tissue probability maps (TPMs) and DARTEL templates^62^, as well as an average T1 anatomical template for visualization purposes^63^. A cohort of 1000 children (500 girls) was selected from the ABCD dataset according to the following procedure. We generated 10,000 random samples with 1000 children (half girls) and selected the one with age and scan site distributions closest to those of the entire cohort. We used SPM Segment to generate the individual’s tissue maps and the TOM8 Toolbox (http://dbm.neuro.uni-jena.de/software/tom) to create the population TPMs and T1 anatomical template, controlling for the effects of age and gender^64^. DARTEL templates were constructed using utilities available in SPM. This involved creating gray (rp1) and white (rp2) matter segments after affine registration followed by DARTEL nonlinear image registration, whereby all selected images were iteratively aligned with a template generated from their own mean, and finally normalized to the MNI space (ICBM template).

### Voxel-based morphometry (VBM)

We implemented VBM analysis to quantify regional gray matter volumes (GMVs) with the CAT12 toolbox (http://dbm.neuro.uni-jena.de/vbm/). The details of VBM analysis have been described in our previous studies^65, 66^. VBM analysis identifies differences in the local composition of brain tissue, accounting for large-scale variation in gross anatomy and location. The analysis includes spatially normalizing individuals’ structural images to the same stereotactic space, segmenting the normalized images into distinct brain tissues, and smoothing the gray matter images. We used the raw images to avoid potential interference with the CAT12 preprocessing pipeline. T1-images were first co-registered to the MNI template using a multiple-stage affine transformation during which the 12 parameters were estimated. Co-registration was performed with a coarse affine registration using mean square differences, followed by a fine affine registration using mutual information. Coefficients of the basis functions that minimized the residual squared difference between the individual image and the template were estimated. Our custom TPMs constructed from 1,000 ABCD children were used in the initial affine transformation. T1 images were then preprocessed with spatial-adaptive non-local means (SANLM) denoising filters^67^ as well as Markov random fields (MRF), corrected for intensity bias field and segmented into cerebrospinal fluid, gray, and white matter^68^. Segmented and the initially registered tissue class maps were normalized using DARTEL^62^, a fast diffeomorphic image registration algorithm of SPM. As a high-dimensional non-linear spatial normalization method, DARTEL generates mathematically consistent inverse spatial transformations. We used our custom DARTEL template in MNI space, as constructed from 1,000 ABCD children, to drive DARTEL normalization. Skull-stripping and final clean-up were performed with default parameters in the CAT12 to remove remaining meninges and correct for volume effects in some regions. In particular, skull-stripping was performed by refining the probability tissue maps of SPM using adaptive probability region-growing, and the final clean-up routine consisted of morphological, distance and smoothing operations after the final segmentation. Normalized GM maps were modulated to obtain the absolute volume of GM tissue corrected for individual brain sizes. Finally, the GM maps were smoothed by convolving with an isotropic Gaussian kernel (FWHM = 8 mm).

Quality check of images was performed visually and quantitatively with tools available in the CAT12 toolbox^69^. One axial slice (z = 0) per subject was plotted and visually checked (option “Display slices”), and outliers were detected by computing the voxel-wise cross-correlation of GM density across subjects (option “Check sample homogeneity”). A total of 47 subjects presented clearly faulty segmentation of brain tissues and were removed from the group analyses. The faulty segmentation likely resulted from poor contrast or artifact of the structural images or abnormal brain shapes. Additionally, for each subject, pairwise correlations were computed voxel-wise between the subject’s GMV and all the other subjects’ GMV. The mean correlation represented how similar the subject’s GMV was to the rest of the sample. Fifty-two subjects with a mean correlation < 0.70, suggesting a higher variance, were also removed.

### Group analyses

In group analyses, we first compared age (in months) between girls and boys. We used two-sample *t* tests to sex differences in ADHD scores and 2-back accuracy rates. We computed correlations between 2-back accuracy rates and ADHD scores in girls and boys separately. Slope tests were used to assess sex differences in the correlations.

For the GMV data, we first examined sex differences in the whole-brain GMV using a two-sample *t* test with age (in months), total intracranial volume (TIV), ADHD score, study site, and scanner model as covariates. We performed a whole-brain linear regression against the ADHD score in girls and boys combined, as well as in girls and boys separately, with age, TIV, study site, and scanner model as covariates. We also performed the same whole-brain linear regression analyses without including TIV as a covariate. The results were evaluated with a voxel *p* < 0.05, corrected for family-wise error of multiple comparisons, based on Gaussian random field theory as implemented in the SPM. Clusters were overlaid on the custom MRI template obtained from the 1,000 ABCD children. Effect sizes were computed using tools available in the CAT12, by approximating Cohen’s *d*^70^ from the *t*-statistics using the expression 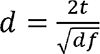 as employed in the study of Kleber et al^71^. The effect sizes of two-sample *t* tests were computed according to the equivalence 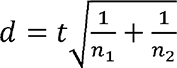 given the sample sizes *n*_1_ and *n*_2_ of the two groups^72^. Customized computations were implemented and verified with the equivalent effect size (Hedge’s g) calculated using the MES toolbox (https://github.com/hhentschke/measures-of-effect-size-toolbox)^73^.

For the regions of interest (ROIs) identified from linear regressions in girls or boys alone, we tested sex differences in the correlation directly with slope tests, with the same set of covariates and showed two-tailed *p* values^74^. Note that the analysis did not represent “double-dipping”, as the slope tests may confirm or refute sex differences^75–79^. This is because the regression maps were identified with a threshold and for example, a cluster showing correlation in boys could show a correlation that just missed the threshold in girls; thus, slope tests were needed to examine whether the correlations were indeed different between the sexes. Sex differences in the extent to which the GMVs correlated with ADHD scores could also be tested in a linear mixed-effects model with an interaction term. We conducted the analyses in SPSS 26.0, with sex, ADHD score, and sex × ADHD score as fixed effects, twin/sibling status as a random effect variable in the model, after controlling for age, race, study site, scanner model, and TIV.

A total of 1,052 children received psychotropic medications (some multiple medications) at the time of the study (psychostimulants: n = 941; antidepressants: n = 176; other psychotropics, including those for epilepsy: n = 25). Children receiving stimulant treatment likely had more significant ADHD symptoms and, as expected, demonstrated significantly higher ADHD scores than unmedicated children (see Results). We conducted a covariance analysis followed by Tukey’s range test to examine the effects of medication on the BG GMVs. Besides, we performed an additional set of whole-brain regression analyses on ADHD scores in medication-naïve children only.

### Heritability of ADHD score and volumetric correlates

Of the current sample, 634 and 2,826 were identified as monozygotic twins (MZ) and dizygotic twins/siblings (DZ), respectively. Thus, we examined whether the ADHD scores, as well as the volumetric correlates, were more significantly related in MZ, as compared to DZ and unrelated pairs (UR), and in DZ as compared to UR. To this end, we computed the Pearson’s correlations each for the ADHD scores, 2-back accuracy rates as well as the volumetric correlates for the MZ and DZ pairs. For the UR, pairs of children were randomly constructed by shuffling and splitting the sample into halves. This procedure was repeated 100 times, and the mean regression lines were computed. For the correlations in MZ and DZ pairs, 95% confidence intervals (CI) were also calculated. We performed slope tests, pairwise, to examine the differences between MZ, DZ and UR. The details of correlation analyses have been described in one of our previous studies^65^. The analyses of the GMVs were done for the clusters combined of those identified with positive and negative correlations, respectively, from all subjects and of the “girls-specific” and “boys-specific” correlates.

For girls and boys combined and separately, we used Mplus 8 to compute the genetic influence (heritability estimates), shared environmental influence, and unique environmental influence^80, 81^ for the ADHD scores and 2-back accuracy rates based on univariate ACE models^82^, with age, race, and study site as covariates. We also estimated univariate ACE models for the volumetric correlates, with scanner model and TIV as additional covariates. Only same-sex DZ pairs were included for the analyses of girls and boys separately. The ACE model decomposes the observed variance into additive genetic factors (A), also known as heritability, shared environmental factors (C), and unique environmental factors (E), in addition to measurement errors^83^. The correlation between the additive genetic variance is fixed to 1.0 for MZ and 0.5 for DZ. The correlation between the shared environmental variance is set to 1.0 for both MZ and the DZ based on the equal environment assumption. The correlation between the non-shared or unique environmental variance is set to zero. The expected variance-covariance matrices within the MZ and DZ are as follows:

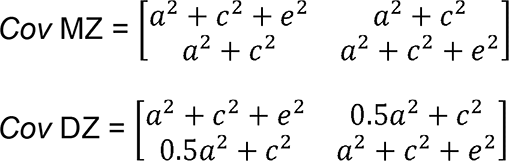

where *a*, *c*, and *e* represent the path coefficients for the A, C, and E factors, respectively^84^. The variance of A, C, and E was estimated using the maximum likelihood method on the variance-covariance matrices and 95% CI of A, C, and E were computed: 95% CI = mean ± 1.96 × standard error, with the assumption that the population standard deviation is a known value. Values of *χ*^2^/*df*, root-mean-square error of approximation (RMSEA), and Tucker-Lewis index (TLI) were used as model fit indices. A *χ*^2^/*df* < 2, an RMSEA < 0.06, or a TLI > 0.95 indicates good fit. Genetic variance (*a*^2^) < 0.30, 0.30-0.60, and > 0.60 is considered low, moderate, and high, respectively^85^. We performed two-sample *t* tests to compare girls and boys in the genetic variance of the ADHD score, sex-specific volumetric correlates, and 2-back accuracy.

Because of the focus on BG, we also estimated the ACE models of the caudate, putamen and pallidum GMVs with the AAL masks for girls and boys combined and separately, with the same set of covariates. Likewise, we performed two-sample *t* tests to examine sex differences.

## Results

### ADHD scores and 2-back accuracy rates

The distributions of ADHD *T* and raw scores of girls and boys are shown in **Supplementary Figure S1**. Girls and boys were different in age (in months), with boys older than girls (mean ± SD: 119.10 ± 7.50 vs. 118.80 ± 7.40 months, *t* = 2.20, *p* = 0.028; two-tailed two-sample *t* test). Boys showed significantly higher ADHD scores than girls (53.61 ± 5.98 vs. 52.71 ± 5.08; *t* = 8.66, *p* < 0.001; two-tailed *t* test with age, race, twin status, and study site as covariates). However, boys and girls showed the same range in ADHD scores (50 to 80) with comparable variability (coefficient of variation: 0.111 for boys and 0.097 for girls).

Boys showed a significantly higher 2-back accuracy rate than girls (75.60 ± 15.09% vs. 73.67 ± 13.98%; *t* = 6.13, *p* < 0.001; two-tailed *t* test with the same covariates). Two-back accuracy rates were significantly and negatively correlated with ADHD scores in boys (*r* = −0.133, *p* < 0.001) and in girls (*r* = −0.116, *p* < 0.001). A slope test showed that boys and girls did not differ significantly in the correlations (*t* = 0.399, *p* = 0.690).

### Sex differences in GMVs

We first examined sex differences in GMV with a two-sample *t* test controlling for age (in months), ADHD score, study site, scanner model, and TIV. As shown in **Supplementary Figure S2**, boys relative to girls showed higher GMVs nearly across the entire brain (*p* < 0.05, FWE corrected). Girls relative to boys showed higher GMVs in relatively small clusters in the left caudate head (x = −6, y = 13, z = 16, 520 voxels, *T* = 6.94), bilateral inferior frontal gyri (x = 33, y = 16, z = 24, 1,407 voxels, *T* = 6.81; x = −34, y = 13, z = 24, 1,180 voxels, *T* = 6.15) and right intraparietal sulcus (x = 35, y = −36, z = 35, 2,377 voxels, *T* = 7.94). The unthresholded map is shown in **Supplementary Figure S5a**.

### GMV correlates of ADHD scores

We examined regional volumetric correlates of ADHD scores in linear regressions for all subjects and girls and boys separately, all with age (in months), study site, scanner model, and TIV as covariates. Brain regions with GMVs in correlation with ADHD scores are shown in Figure 1 and the clusters are summarized in **Supplementary Table S1** (*p* < 0.05, FWE corrected). Overall, in boys and girls combined, ADHD scores were positively correlated with a single cluster in the area of the hypothalamus (x = −5, y = −13, z = −9, voxel *Z* = 6.88, 689 voxels). The inset of Figure 1 shows the cluster overlaid on a hypothalamus mask^86, 87^. In contrast, higher ADHD scores were associated with the reduction of GMVs in an extensive array of brain regions, including bilateral caudate and putamen, orbitofrontal cortex (OFC), visual cortex, somatomotor cortex, and temporal cortex. The unthresholded map is shown in **Supplementary Figure S5b**.

**Figure 1.**
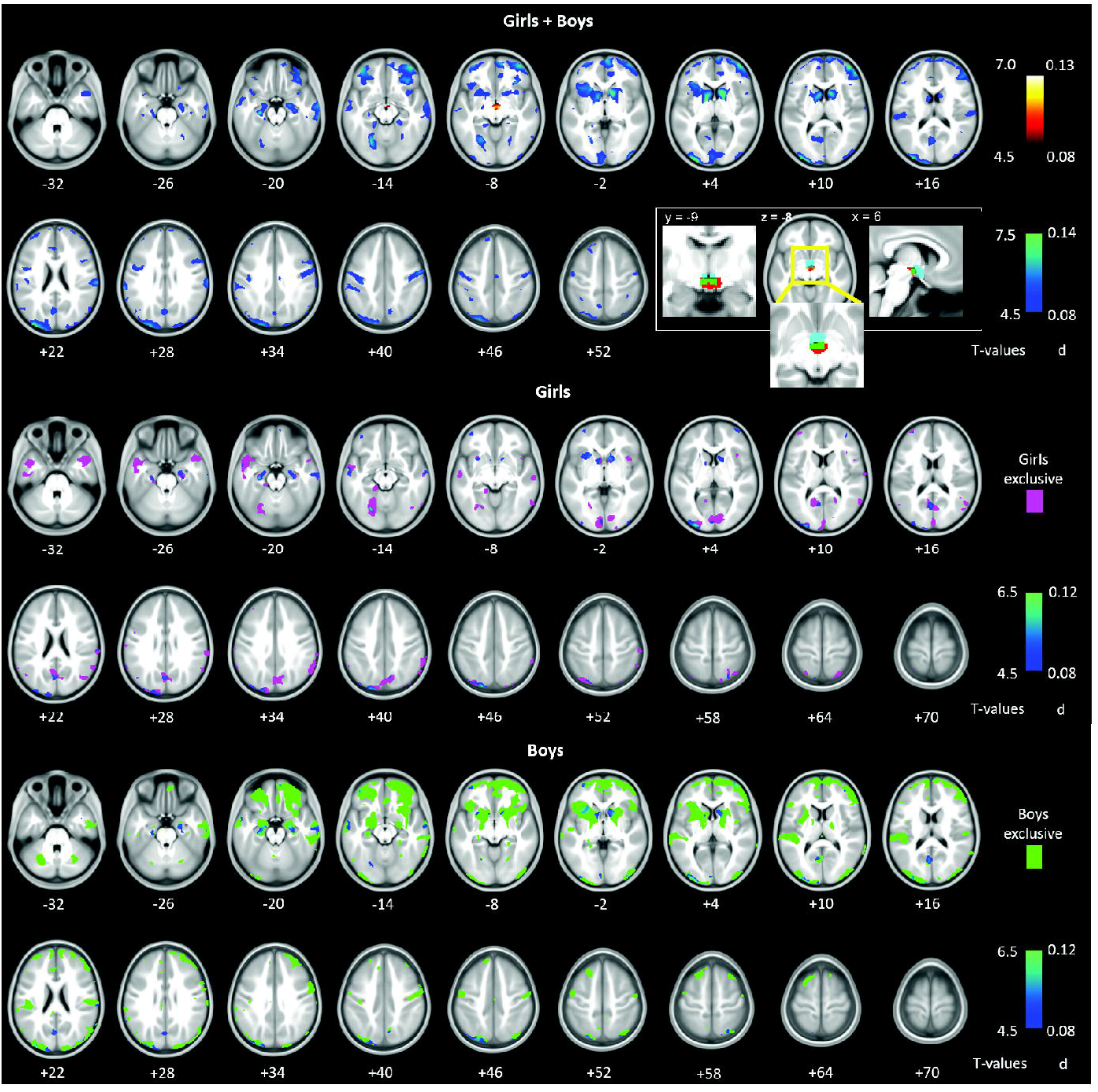
GMV correlates of ADHD scores for girls and boys combined (top panels), and for girls (middle) and boys (bottom), separately. Whole-brain regression, *p* < 0.05, FWE corrected. Color bars show voxel *T* values (left, cool colors to indicate negative correlation) and the corresponding effect sizes (right, Cohen’s *d*). A single cluster in the area of the hypothalamus showed a positive correlation with ADHD scores in girls and boys combined; the inset shows the cluster overlaid on a hypothalamus mask (cyan) with overlapping voxels highlighted in dark green. For the clusters identified of girls and boys separately, we used exclusive masking to highlight those that appeared specific to girls (pink) and to boys (light green). Overall, boys relative to girls showed more widespread and significant GMV reductions in relation to ADHD scores.

Except for the hypothalamus, 2-back accuracy rate (%) were positively correlated with the volumetric correlates identified of boys or girls or of all three BG subregions (*r*’s ranging from 0.066 to 0.190, all *p*’s < 0.001) in linear regressions with the same set of covariates. None of these correlations showed a significant sex difference in slope. These results are shown in **Supplementary Table S2**.

In multiple regressions for boys and girls separately, overall boys relative to girls appeared to show more significant reductions in GMV in link with higher ADHD scores, broadly in the frontal (particularly orbitofrontal) and parietal cortical regions as well as subcortical structures, such as the thalamus. In contrast, girls relative to boys appeared to demonstrate more significant GMV reductions in association with higher ADHD scores in the bilateral inferior/middle temporal cortex, frontopolar cortex, medial occipital cortex, and the precuneus. No voxels showed GMVs in positive correlations with ADHD scores in girls or boys alone. These findings are shown in Figure 1, where we highlighted the voxels that appeared to be specific to girls by masking the findings with boys’ clusters, and vice versa.

As some morphometric studies of ADHD also showed imaging findings without specifically accounting for TIV. Thus, we performed additional analyses without including TIV as a covariate and showed the results evaluated at *p* < 0.05 FWE corrected in **Supplementary Figure S3**. Overall, the findings are similar though more prominent. For instance, in girls and boys combined, the ADHD trait was associated with higher GMV of the hypothalamus and lower GMVs of the OFC and BG. The unthresholded map is shown in **Supplementary Figure S5c**.

### Sex-specific GMV correlates of ADHD scores

We extracted the GMV estimates of the clusters identified from whole-brain analyses and performed slope tests to examine whether there were sex differences in the linear regressions of the GMVs against ADHD scores with age, race, twin status, study site, scanner model, and TIV as covariates. Slope tests did not reveal significant sex differences in the slopes of the regressions of hypothalamus GMVs vs. ADHD scores (*t* = 0.250, *p* = 0.803) (Figure 2A). We combined the clusters identified each for girls and boys as “girls’ or boys’ joint ROIs”. We did the same for those “sex-specific” voxels as “girls’ or boys’ specific joint ROIs”. The results showed that the association between GMVs and ADHD scores was more negative in boys (*r* = −0.112, *p* < 0.001) than in girls (*r* = −0.072, *p* < 0.001) for boys’ joint ROIs, with a significant slope difference (*t* = 2.788, *p* = 0.005; corrected *p* = 0.05/8 = 0.00625; Figure 2B). Likewise, the association between GMVs and ADHD scores was more negative in boys (*r* = −0.112, *p* < 0.001) than in girls (*r* = −0.068, *p* < 0.001) for boys’ specific joint ROIs, with a significant slope difference (*t* = 2.99, *p* = 0.003; corrected *p* = 0.05/8 = 0.00625; Figure 2D). None of the girls’ joint ROIs showed a correlation with ADHD scores significantly different between girls and boys in slope tests (both *p’s* ≥ 0.248, Figure 2C and 2E).

**Figure 2.**
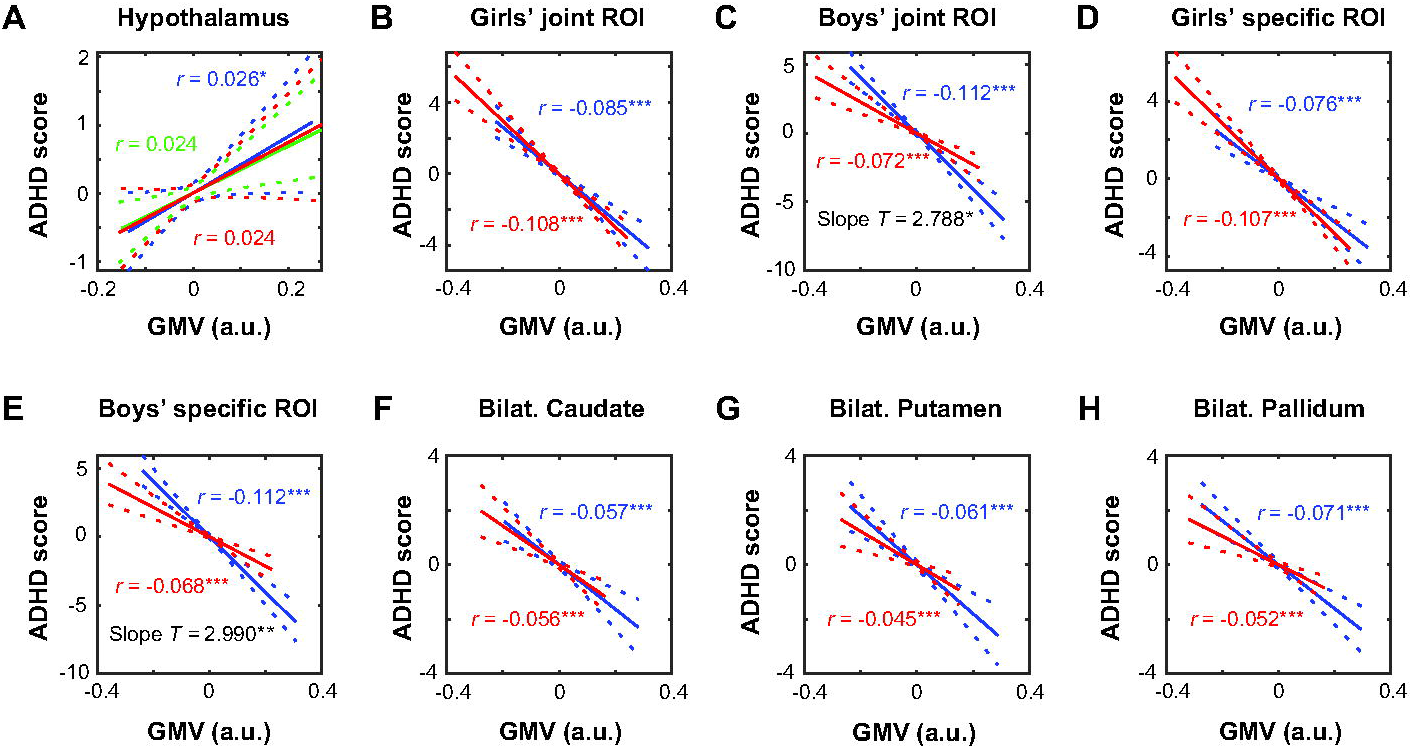
Linear correlations between ADHD scores and GMVs for (**A**) hypothalamus, (**B**) girls’ joint ROIs, (**C**) boys’ joint ROIs, (**D**) girls’ specific ROIs, (**E**) boys’ specific ROIs, (**F**) bilateral caudate, (**G**) bilateral putamen, and (**H**) bilateral pallidum, plotted separately for girls (in red) and boys (in blue), with age, race, twin status, study site, scanner model, and TIV as covariates. Linear correlation for girls and boys combined is also shown for hypothalamus (in green). Only boys’ joint and specific joint ROIs showed GMVs in negative correlation with ADHD score more significantly in boys than in girls (*p’s* < 0.05/8 = 0.00625). Note that the residual scores after controlling for covariates are shown here. Dashed lines represent 95% confidence intervals of the mean regressions (solid lines). a.u. = arbitrary unit of GMV estimates; Bilat. = Bilateral. \*\*\**p* < 0.001, \*\**p* < 0.005, and *\*p* < 0.05. See **Supplementary Table S3** for statistics.

We also examined sex differences specifically for the BG subregions, using the AAL masks, and the results showed girls and boys did not differ in the correlations of GMVs with ADHD scores for bilateral caudate, putamen, or pallidum (all *p’s* ≥ 0.159, slope tests, Figure 2F, 2G**, and** 2H).

**Supplementary Table S3** summarizes the statistics of individual regressions in girls and boys separately and slope tests on sex differences. Linear mixed-effects models showed that the interaction term was significant for boys’ specific joint ROIs (*p* = 0.034), consistent with the slope tests, but not any other ROIs (all *p’s* ≥ 0.057). Note that none of the results on sex differences would be significant with correction for multiple comparisons. The results are summarized in **Supplementary Table S4**.

### The effects of medications

A total of 941 (8.2%) children received psychostimulant medications, who relative to unmedicated children showed more significant ADHD symptoms [ADHD score (mean ± SD): 60.70 ± 7.96 vs. 52.51 ± 4.78; *F* = 2,164.32; *p* < 0.001, covariance analysis with age, sex, race, study site, and twin status as covariates]. The GMVs (mean ± SD) of the caudate were 0.398 ± 0.046 and 0.393 ± 0.049, of the putamen were 0.482 ± 0.049 and 0.481 ± 0.050, and of the pallidum were 0.531 ± 0.060 and 0.524 ± 0.058 for unmedicated and stimulant-treated children, respectively. Covariance analysis with age, sex, race, study site, twin status, scanner model, and TIV as covariates showed a significant effect of psychostimulant for the GMVs of bilateral caudate (*F* = 41.63, *p* < 0.001), putamen (*F* = 51.32, *p* < 0.001), and pallidum (*F* = 72.94, *p* < 0.001). Thus, psychostimulant-treated children showed higher ADHD scores and relatively smaller BG GMVs as compared to medication-naïve children.

In an additional analysis, we excluded all medicated children and performed the same set of regression analyses on ADHD scores (n = 10,450; 5,174 girls). The results showed largely similar volumetric correlates of ADHD scores, though diminished in effect sizes, likely because of the exclusion of the children with more significant ADHD traits (**Supplementary Figure S4** and **Table S5,** *p* < 0.05 FWE corrected). The unthresholded map is shown in **Supplementary Figure S5d**.

### Heritability of ADHD score and volumetric correlates

We computed the Pearson’s correlations between MZ pairs, between DZ pairs and between UR pairs in ADHD scores and volumetric correlates for all and for girls and boys separately. The results are shown in Figure 3. Overall, for both ADHD scores and the volumetric correlates, the correlations were significantly stronger in MZ pairs than in DZ and UR pairs as well as in DZ pairs than in UR pairs. **Supplementary Tables S6 and S7** summarize the statistics of individual sets of correlation and of slope tests for ADHD scores and GMV correlates, respectively.

**Figure 3.**
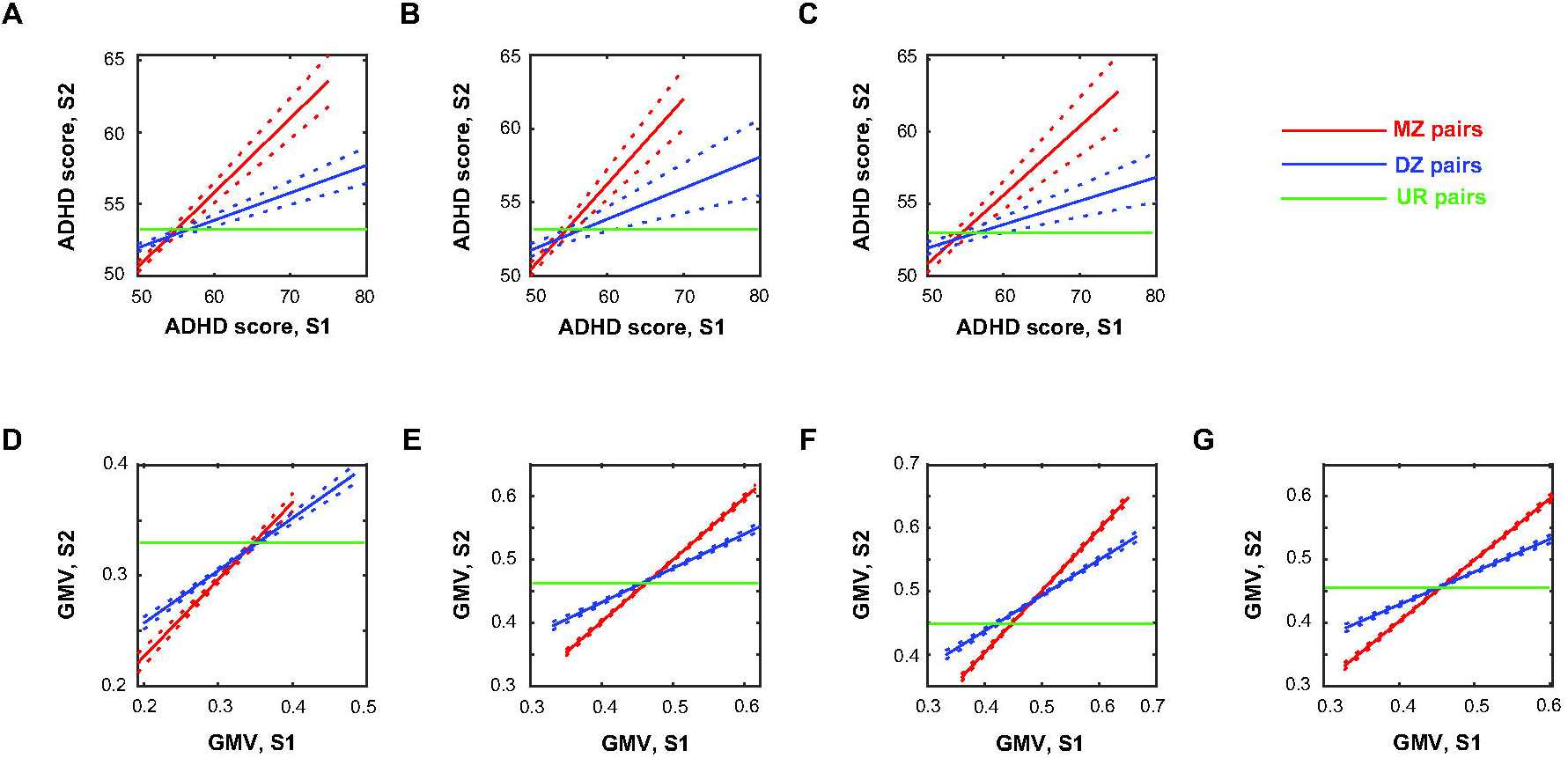
Pearson’s correlations between monozygotic (MZ) pairs, between dizygotic/siblings (DZ) pairs and between unrelated (UR) pairs in ADHD score for (**A**) all, (**B**) girls, and (**C**) boys; and in GMV correlates in (**D**) positive and (**E**) negative correlations with ADHD scores for all, as well as (**F**) “girls-specific” and (**G**) “boys-specific” GMV correlates for all. S1 and S2: subject 1 and 2 of the pair. The *r* and *p* values of the correlations and the results of slope tests are shown in **Supplementary Tables S6 and S7**, respectively.

Table 1 shows the heritability estimates and shared and non-shared environmental effects of ADHD scores and 2-back accuracy rates with age, sex (only for “all”), race, and study site as covariates as well as volumetric correlates with scanner model and TIV as additional covariates in girls and boys combined and separately. The ACE models showed good model fit for the great majority of measures (**Supplementary Table S8**). For girls and boys combined and separately, the ADHD scores (*a^2^* = 0.54- 0.59) were moderately heritable, GMV correlates (*a^2^* = 0.60-0.79) were highly heritable, and 2-back accuracy rates were near moderately heritable (*a^2^* = 0.30-0.39). The shared-environmental effects on the measures were negligible or weak (*c*^2^ = 0.00-0.19). Two-sample *t* tests showed no significant sex differences in the heritability estimates of ADHD scores, volumetric correlates, or 2-back accuracy rates (all *p’s* ≥ 0.764).

**Table 1.** Genetic, shared environmental, and non-shared environmental effects of ADHD scores, GMV correlates, and 2-back%.

| *Variable* | $a^2$ (A) | $c^2$ (C) | $e^2$ (E) |
| --- | --- | --- | --- |
| ADHD score (all) | 0.59 [0.53, 0.66] | 0.00 [0.00, 0.00] | 0.41 [0.34, 0.48] |
| ADHD score (girls) | 0.54 [0.35, 0.74] | 0.00 [0.00, 0.00] | 0.46 [0.26, 0.65] |
| ADHD score (boys) | 0.56 [0.46, 0.66] | 0.00 [0.00, 0.00] | 0.44 [0.34, 0.54] |
| GMV positive correlate (all, HT) | 0.60 [0.45, 0.74] | 0.04 [-0.06, 0.15] | 0.36 [0.30, 0.42] |
| GMV negative correlate (all) | 0.77 [0.68, 0.86] | 0.07 [-0.02, 0.15] | 0.16 [0.14, 0.19] |
| GMV correlate (“girls specific”) | 0.66 [0.56, 0.74] | 0.19 [0.12, 0.26] | 0.15 [0.13, 0.18] |
| GMV correlate (“boys specific”) | 0.79 [0.69, 0.89] | 0.03 [-0.05, 0.12] | 0.18 [0.15, 0.21] |
| 2-back% (all) | 0.30 [0.07, 0.53] | 0.08 [-0.07, 0.23] | 0.62 [0.52, 0.72] |
| 2-back% (girls) | 0.33 [-0.02, 0.68] | 0.02 [-0.21, 0.26] | 0.65 [0.49, 0.80] |
| 2-back% (boys) | 0.39 [0.05, 0.72] | 0.03 [0.20, 0.27] | 0.58 [0.44, 0.72] |

*Note:* HT: hypothalamus; $a^2$ : proportion of variance due to additive genetic effects (A); $c^2$ : proportion of variance due to shared environmental effects (C); $e^2$ : proportion of variance due to non-shared environmental effects (E); 95% confidence intervals are presented in square brackets. Age, sex (only for “all”), study site, race, and (for GMVs) scanner model and TIV were also included as covariates for GMVs. Only 9,982 children, with 500 MZ and 2,208 DZ had N-back task data.

The model fits and ACE estimation for the GMVs of BG subregions are shown in **Supplementary Table S9a** and **S9b**, respectively. The ACE models showed good model fit for the great majority of these volumetric measures. The heritability of caudate, putamen and pallidum volumes were in the range of 0.8 to 0.9, slightly higher than those reported in an earlier study of family-based cohorts^88^. Two-sample *t* tests showed that there were no significant differences in the heritability of caudate, putamen, or pallidum GMVs between girls and boys (all *p’s* ≥ 0.936).

## Discussion

We observed negative correlations between the ADHD scores and GMVs in a wide array of brain regions in both boys and girls, in line with previous findings^17, 89–91^. More significant and broadly distributed in boys than in girls, the volumetric reductions clearly involved bilateral caudate and putamen. As part of the cognitive control network^92–94^, the BG play important roles in executive functions such as attention, memory, response inhibition, and set-shifting, known to be impaired in ADHD^95, 96^. Indeed, we showed that ADHD scores and bilateral caudate GMVs were negatively and postively, respectively, correlated with 2-back accuracy in both girls and boys. Both ADHD traits and the volumetric markers are heritable. The heritability of ADHD scores, volumetric markers, and working memory were similar between the sexes. Together, the current findings confirm BG volumetric reduction in children with ADHD traits, overall more significant GMV reductions in boys than in girls, and the roles of caudate GMV reduction in working memory dysfunction. We have also observed a cluster in the hypothalamus that showed GMV in positive correlation with ADHD score across all subjects but not in boys or girls alone. With future releases of the ABCD data, it would be of tremendous interest and translational significance to investigate how these structural brain markers predict the development of full-blown ADHD and comorbidities in these children. We discussed the main findings below.

### Volumetric correlates of ADHD traits

We observed a cluster in the hypothalamus that showed GMV in positive correlation with ADHD scores across all subjects but not in boys or in girls alone examined at the same threshold. The hypothalamus is known for its role in motivated behavior and stress response. Studies have reported altered activity of the hypothalamus-pituitary axis, as evidenced in the abnormal diurnal rhythm of cortisol secretion and cortisol response to stress, in children with ADHD^97–99^. However, it remains unclear whether the neuroendocrine abnormalities reflect experienced stress or the pathophysiological process of ADHD per se^97, 100^. A recent imaging study implicated hypothalamic dysfunction during post-error slowing in the stop signal task, a deficit correlated with the inattention score, in children with ADHD^101^. In broad consistency, earlier volumetric studies have associated smaller hypothalamus GMV with behavioral inhibition traits in children^65^ and young adults^102^. The functional implication of enlarged hypothalamus in children with higher ADHD traits, if replicated, remains to be investigated.

Most notable among the cortical regions showing significant GMV reductions in link with ADHD scores in boys are bilateral, including both medial and lateral, OFC. The OFC and orbito-frontal-limbic circuit have been implicated in emotional impulsivity^27^, and volumetric deficits of the OFC appear consistent with affective dysfunction in some patients with ADHD^89^. On the other hand, few extant VBM studies reported OFC GMV reduction in ADHD^27, 52, 103^. Children in the ABCD cohort were of 9 to 10 years of age, younger than the subject population reported in most of the earlier studies, which have suggested age-related changes in the volumetric correlates of ADHD^19, 104^. For instance, machine learning demonstrated that although structural MRI data can significantly separate ADHD from control participants for both children and adults, prediction performance and effect sizes were better for the child than the adult samples^105^. It thus remains to be seen whether OFC volumetric deficits may perhaps normalize with age but would continue to manifest in individuals with higher emotional impulsivity.

In girls and boys combined as well as in boys alone, the occipital cortex showed diminished GMV in relation to ADHD traits, consistent with earlier reports of occipital cortical GMV reduction in children with ADHD with a mean age of around 10 years^106, 107^ and in adolescents with ADHD^24^. Other studies showed that the baseline GMV of the occipital cortex predicted inattention symptoms in a 2-year follow-up and was associated with the genetic risk for ADHD^108^. The GMV of the occipital cortex may predict stimulant treatment response in ADHD^109^. Another study of college students associated greater impulsivity scores to higher volumes in the occipital cortex specifically in the region that represents the peripheral visual field^110^. Together, prior literature provides evidence implicating occipital cortical GMV in ADHD traits, in accord with the visual cortical dysfunction in link with attention deficits in ADHD^111^.

### Sex differences in cerebral volumetric reductions in ADHD

The volumetric changes associated with ADHD scores are more remarkable and broadly distributed in boys, consistent with a higher incidence and greater severity of ADHD in boys than in girls^23, 32, 33, 112^. In contrast, although girls relative to boys appeared to show more significant deficits in the inferior/middle temporal cortex and the precuneus, the differences did not show significantly in slope tests. In boys relative to girls, more widespread deficits were found in the OFC, frontoparietal regions, including the ACC, and subcortical structures such as the thalamus and BG, which are the core structures of the cognitive control network^113, 114^. These sex differences are confirmed by slope tests but not by the interaction effects with correction for multiple comparisons, and the slope tests restricted to the BG nuclei did not demonstrate significant sex differences^115–117^. These findings are broadly consistent with those reported from the ENIGMA studies of ADHD^118^ and those implicating a link between caudate nucleus and deficits of executive function in ADHD^27, 111^. On the other hand, although the findings together are commensurate with the proposition of frontal-striatal network dysfunction in ADHD, evidence also suggests the roles of other cortical structures, including the occipital cortical regions, in the etiological processes of attention deficits in ADHD^111^. Our findings enrich the theory by identifying the structural deficits of these regions in ADHD and provide a potential explanation of more significant ADHD symptoms in boys. Note that boys and girls did not demonstrate significant differences in the correlations of ADHD scores or volumetric correlates with 2-back accuracy, or in the heritability of the volumetric markers, ADHD score, or N-back performance. We speculate that these volumetric correlates may more broadly represent neural markers of sex differences in clinical manifestations, including greater impulsivity in boys than in girls with ADHD^119–121^, which should be explored further. Studies investigating the heritability, particularly those involving the computation of polygenic risk scores, of both neural and behavioral markers would help addressing the inter-relationships between gene, brain, and behavior in ADHD.

### Medications and volumetric reductions

As expected, children medicated with stimulants showed higher ADHD scores than those who were medication naïve. We demonstrated in medication-naïve children similar albeit slightly weaker volumetric reductions in association with ADHD traits. In particular, the volumetric reductions in the putamen and pallidum were no longer significant, suggesting that the lentiform nucleus may play a unique role in the pathogenesis of more significant forms of ADHD. Indeed, studies focusing on subcortical structures have typically reported putamen volumetric deficits in ADHD^27, 122^. Further, lesions of the putamen have been associated with the development of secondary ADHD^123, 124^. In contrast, the GMV reduction of the precuneus remains strongly associated with ADHD scores in medication-naïve girls, suggesting potentially distinct volumetric markers of ADHD traits in the female sex.

### Heritability of ADHD traits, volumetric markers, and working memory

ADHD traits in children were near highly heritable (*a*^2^ = 0.59, all subjects), suggesting that variance in the ADHD symptoms can largely be accounted for by genetic factors^125–127^. In accord with previous findings^42^, the volumetric correlates were also highly heritable, with *a*^2^ = 0.60 for the hypothalamus and *a*^2^ ∼ 0.70 for the “negative” correlates. On the other hand, studies have also supported the roles of environmental factors in the development of ADHD^128, 129^. For instance, ADHD has been associated with low socio-economic status^130^. In particular, with *e^2^* at 0.41 across all subjects, 0.46 in girls, and 0.44 in boys, unique environmental factors modestly explained the variance in ADHD symptoms, in accord with previous reports^131^. An earlier longitudinal study showed that as the ADHD symptoms evolved from childhood to adolescence, the unique environmental influences may result from a socialization process dominated by interpersonal relationships with parents, teachers and peers^36^.

The GMVs of the BG subregions (caudate, putamen and globus pallidum) were strongly influenced by genetics (*a*^2^ = 0.79-0.90; all subjects), largely in line with earlier reports that genetic variation accounted for 43-85% of the variance in the BG subregional GMVs in children^132–134^. Other studies with ∼200 twin pairs showed comparable heritability for BG GMVs in young and old adults (*a*^2^ = 0.65-0.88)^135–137^. Thus, the heritability of BG GMVs appears to be stable across age. Two-back accuracy rate was moderately heritable (*a*^2^ = 0.30; all subjects), though not to the extent observed in young adults (*a*^2^ = 0.73; 60 twin pairs)^138^. The trend of increased heritability for working memory across age has also been observed for other cognitive functions^85, 139^.

We did not observe sex differences in the heritability of ADHD traits, volumetric markers, or working memory, in accord with an earlier work^140^. Another study likewise found no evidence for sex differences in the genetic influences on externalizing disorders, including ADHD, conduct disorder, and oppositional defiant disorder in adolescents^141^. Given that ADHD appears to be more prevalent in the male sex, the sex difference may manifest in other behavioral or structural or functional neural markers of ADHD. Identifying the sex-related markers would help in investigating sex-specific genetic polymorphisms of ADHD^140, 142^.

### Limitations of the study and conclusions

A number of limitations need to be considered. First, ADHD is known to be comorbid with many behavioral conditions, including depressive^143^ and conduct disorders^144^. We did not control for these and many other clinical variables in data analyses; thus, the impact of comorbidities on the current findings remains to be clarified, as demonstrated in the ENIGMA studies of ADHD^145–147^. Second, the correlations between ADHD scores and GMVs are relatively weak, which may reflect the non-clinical populations of the ABCD cohort. On the other hand, it is worth noting that the effect sizes were within the range reported of volumetric markers of ADHD cases^19^. Third, pubertal maturation may contribute to sex differences in brain development^148^. The ABCD study includes pubertal survey and provides an opportunity for future work to investigate the effects pubertal age on brain development^149^. Fourth, we acknowledge the limitations of twin design analyses, including the difficulty in detecting shared environmental effects and inflated estimates of heritability^84, 150^. Fifth, these are cross-sectional findings; with release of more and more follow-up data, investigators would be able to examine how the volumetric deficits evolve along with clinical manifestations of ADHD into late adolescence. Finally, whereas the current work focuses on quantifying GMVs, previous studies have investigated other morphometric measures, including cortical thickness and surface area, to better understand cerebral structural alterations in ADHD^32, 33, 54^.

To conclude, we confirmed cerebral, including basal ganglia, volumetric deficits of ADHD traits in children and potential sex differences in these structural alterations in relation to ADHD traits. ADHD traits and the volumetric markers are highly heritable with no evidence of sex differences in heritability. These findings would inform future research of the neural markers of ADHD and of the clinical trajectories and efficacy of treatment as the ABCD children are followed through late adolescence.

## Acknowledgements

This study is supported by NIH grant DA023248, DA044749, DA045189 and DA051922. The NIH is otherwise not responsible for the conceptualization of the study, data collection and analysis, or in the decision in publishing the results. MATLAB and other codes are available on GitHub (https://github.com/jaimeide/LiLabYale/tree/main/LiChenIde_SR2022).

## Author contributions

All authors contributed to conceptualization of the study, data analysis and interpretation, literature review, and writing of the manuscript.

## Additional Information

### Data availability, Ethics declarations, and Consent to participate

We have obtained permission from the ABCD to use the Open and Restricted Access data for the current study. The ABCD data is publicly available at https://abcdstudy.org/. The ABCD data repository grows and changes over time. The data used in this report (NDA project ID 2573, Release 2.0) came from http://dx.doi.org/10.15154/1503209. All recruitment procedures and informed consent forms, including consent to share de-identified data, were approved by the Institutional Review Board of the University of California San Diego with study number 160091. Informed consent was obtained from all individual participants included in the study.

## References

1 Fayyad, J. et al. Cross-national prevalence and correlates of adult attention-deficit hyperactivity disorder. British Journal of Psychiatry 190, 402–409, doi:10.1192/bjp.bp.106.034389 (2007).

2 Thomas, R., Sanders, S., Doust, J., Beller, E. & Glasziou, P. Prevalence of attention-deficit/hyperactivity disorder: a systematic review and meta-analysis. Pediatrics 135, e994–e1001 (2015).

3 Danielson, M. L. et al. Prevalence of parent-reported ADHD diagnosis and associated treatment among US children and adolescents, 2016. Journal of Clinical Child & Adolescent Psychology 47, 199-212 (2018).

4 Willcutt, E. G., Doyle, A. E., Nigg, J. T., Faraone, S. V. & Pennington, B. F. Validity of the executive function theory of attention-deficit/hyperactivity disorder: a meta-analytic review. Biological psychiatry 57, 1336–1346 (2005).

5 Johnson, K. A., Wiersema, J. R. & Kuntsi, J. What would Karl Popper say? Are current psychological theories of ADHD falsifiable? Behavioral and Brain Functions 5, 15 (2009).

6 Rapport, M. D. et al. Working memory deficits in boys with attention-deficit/hyperactivity disorder (ADHD): The contribution of central executive and subsystem processes. Journal of abnormal child psychology 36, 825–837 (2008).

7 Kofler, M. J. et al. Working memory deficits and social problems in children with ADHD. Journal of abnormal child psychology 39, 805–817 (2011).

8 Ehlis, A.-C., Bähne, C. G., Jacob, C. P., Herrmann, M. J. & Fallgatter, A. J. Reduced lateral prefrontal activation in adult patients with attention-deficit/hyperactivity disorder (ADHD) during a working memory task: a functional near-infrared spectroscopy (fNIRS) study. Journal of psychiatric research 42, 1060–1067 (2008).

9 Massat, I. et al. Working memory-related functional brain patterns in never medicated children with ADHD. PloS one 7, e49392 (2012).

10 Jaeggi, S. M. et al. The relationship between n-back performance and matrix reasoning—implications for training and transfer. Intelligence 38, 625–635 (2010).

11 Li, G. et al. Neural correlates of individual variation in two-back working memory and the relationship with fluid intelligence. Sci Rep 11, 9980, doi:10.1038/s41598-021-89433-8 (2021).

12 Chen, Y. et al. Testing a Cognitive Control Model of Human Intelligence. Sci Rep 9, 2898, doi:10.1038/s41598-019-39685-2 (2019).

13 Cubillo, A., Halari, R., Smith, A., Taylor, E. & Rubia, K. A review of fronto-striatal and fronto-cortical brain abnormalities in children and adults with Attention Deficit Hyperactivity Disorder (ADHD) and new evidence for dysfunction in adults with ADHD during motivation and attention. cortex 48, 194–215 (2012).

14 Rommelse, N., Buitelaar, J. K. & Hartman, C. A. Structural brain imaging correlates of ASD and ADHD across the lifespan: a hypothesis-generating review on developmental ASD–ADHD subtypes. Journal of Neural Transmission 124, 259–271 (2017).

15 Valera, E. M., Faraone, S. V., Murray, K. E. & Seidman, L. J. Meta-analysis of structural imaging findings in attention-deficit/hyperactivity disorder. Biological psychiatry 61, 1361–1369 (2007).

16 Ellison-Wright, I., Ellison-Wright, Z. & Bullmore, E. Structural brain change in Attention Deficit Hyperactivity Disorder identified by meta-analysis. BMC Psychiatry 8, doi:10.1186/1471-244x-8-51 (2008).

17 Nakao, T., Radua, J., Rubia, K. & Mataix-Cols, D. Gray matter volume abnormalities in ADHD: voxel-based meta-analysis exploring the effects of age and stimulant medication. American Journal of Psychiatry 168, 1154–1163 (2011).

18 Frodl, T. & Skokauskas, N. Meta-analysis of structural MRI studies in children and adults with attention deficit hyperactivity disorder indicates treatment effects. Acta Psychiatrica Scandinavica 125, 114–126, doi:10.1111/j.1600-0447.2011.01786.x (2012).

19 Hoogman, M. et al. Subcortical brain volume differences in participants with attention deficit hyperactivity disorder in children and adults: a cross-sectional mega-analysis. The Lancet Psychiatry 4, 310–319 (2017).

20 Stoodley, C. J. Distinct regions of the cerebellum show gray matter decreases in autism, ADHD, and developmental dyslexia. Frontiers in systems neuroscience 8, 92–92, doi:10.3389/fnsys.2014.00092 (2014).

21 Lim, L. et al. Disorder-specific grey matter deficits in attention deficit hyperactivity disorder relative to autism spectrum disorder. Psychological Medicine 45, 965–976, doi:10.1017/s0033291714001974 (2014).

22 Makris, N. et al. Toward Defining the Neural Substrates of ADHD. Journal of Attention Disorders 19, 944–953, doi:10.1177/1087054713506041 (2013).

23 Villemonteix, T. et al. Grey matter volume differences associated with gender in children with attention-deficit/hyperactivity disorder: a voxel-based morphometry study. Developmental cognitive neuroscience 14, 32–37 (2015).

24 Bonath, B., Tegelbeckers, J., Wilke, M., Flechtner, H.-H. & Krauel, K. Regional gray matter volume differences between adolescents with ADHD and typically developing controls: further evidence for anterior cingulate involvement. Journal of attention disorders 22, 627–638 (2018).

25 Bralten, J. et al. Voxel-based morphometry analysis reveals frontal brain differences in participants with ADHD and their unaffected siblings. Journal of psychiatry & neuroscience: JPN 41, 272 (2016).

26 Kumar, U., Arya, A. & Agarwal, V. Neural alterations in ADHD children as indicated by voxel-based cortical thickness and morphometry analysis. Brain and Development 39, 403–410, doi:10.1016/j.braindev.2016.12.002 (2017).

27 Jagger-Rickels, A. C., Kibby, M. Y. & Constance, J. M. Global gray matter morphometry differences between children with reading disability, ADHD, and comorbid reading disability/ADHD. Brain and Language 185, 54–66, doi:10.1016/j.bandl.2018.08.004 (2018).

28 Lukito, S. et al. Comparative meta-analyses of brain structural and functional abnormalities during cognitive control in attention-deficit/hyperactivity disorder and autism spectrum disorder. Psychological Medicine 50, 894–919, doi:10.1017/s0033291720000574 (2020).

29 Zhao, Y. et al. Aberrant gray matter volumes and functional connectivity in adolescent patients with ADHD. Journal of Magnetic Resonance Imaging 51, 719–726, doi:10.1002/jmri.26854 (2019).

30 Rucklidge, J. J. Gender differences in attention-deficit/hyperactivity disorder. Psychiatric Clinics 33, 357–373 (2010).

31 Greven, C. U., Rijsdijk, F. V. & Plomin, R. A twin study of ADHD symptoms in early adolescence: hyperactivity-impulsivity and inattentiveness show substantial genetic overlap but also genetic specificity. Journal of abnormal child psychology 39, 265–275 (2011).

32 Qiu, A. et al. Basal ganglia volume and shape in children with attention deficit hyperactivity disorder. American Journal of Psychiatry 166, 74–82 (2009).

33 Onnink, A. M. H. et al. Brain alterations in adult ADHD: effects of gender, treatment and comorbid depression. European Neuropsychopharmacology 24, 397–409 (2014).

34 Villemonteix, T. et al. Grey matter volumes in treatment naïve vs. chronically treated children with attention deficit/hyperactivity disorder: a combined approach. European Neuropsychopharmacology 25, 1118–1127, doi:10.1016/j.euroneuro.2015.04.015 (2015).

35 Faraone, S. V. et al. Molecular genetics of attention-deficit/hyperactivity disorder. Biological psychiatry 57, 1313–1323 (2005).

36 Larsson, J.-O., Larsson, H. & Lichtenstein, P. Genetic and environmental contributions to stability and change of ADHD symptoms between 8 and 13 years of age: a longitudinal twin study. Journal of the American Academy of Child & Adolescent Psychiatry 43, 1267–1275 (2004).

37 Knopik, V. S. et al. Contributions of parental alcoholism, prenatal substance exposure, and genetic transmission to child ADHD risk: a female twin study. Psychological medicine 35, 625–635 (2005).

38 Polderman, T. J. et al. Across the continuum of attention skills: a twin study of the SWAN ADHD rating scale. Journal of Child Psychology and Psychiatry 48, 1080–1087 (2007).

39 Larsson, H., Chang, Z., D’Onofrio, B. M. & Lichtenstein, P. The heritability of clinically diagnosed attention-deficit/hyperactivity disorder across the life span. Psychological medicine 44, 2223 (2014).

40 Chen, Q. et al. Familial aggregation of attention-deficit/hyperactivity disorder. Journal of Child Psychology and Psychiatry 58, 231–239 (2017).

41 Eilertsen, E. M. et al. Development of ADHD symptoms in preschool children: Genetic and environmental contributions. Development and psychopathology 31, 1299–1305 (2019).

42 Durston, S. et al. Magnetic resonance imaging of boys with attention-deficit/hyperactivity disorder and their unaffected siblings. Journal of the American Academy of Child & Adolescent Psychiatry 43, 332–340 (2004).

43 Pironti, V. A. et al. Neuroanatomical abnormalities and cognitive impairments are shared by adults with attention-deficit/hyperactivity disorder and their unaffected first-degree relatives. Biological Psychiatry 76, 639–647 (2014).

44 Greven, C. U. et al. Developmentally stable whole-brain volume reductions and developmentally sensitive caudate and putamen volume alterations in those with attention-deficit/hyperactivity disorder and their unaffected siblings. JAMA psychiatry 72, 490–499 (2015).

45 Khadka, S. et al. Multivariate imaging genetics study of MRI gray matter volume and SNPs reveals biological pathways correlated with brain structural differences in attention deficit hyperactivity disorder. Frontiers in Psychiatry 7, 128 (2016).

46 Xu, B. et al. Impact of a common genetic variation associated with putamen volume on neural mechanisms of attention-deficit/hyperactivity disorder. Journal of the American Academy of Child & Adolescent Psychiatry 56, 436–444. e434 (2017).

47 Luo, X. et al. KTN1 variants and risk for attention deficit hyperactivity disorder. American Journal of Medical Genetics Part B: Neuropsychiatric Genetics 183, 234–244 (2020).

48 Ohi, K. et al. Genetic correlations between subcortical brain volumes and psychiatric disorders. Br J Psychiatry 216, 280–283, doi:10.1192/bjp.2019.277 (2020).

49 Klein, M. et al. Genetic Markers of ADHD-Related Variations in Intracranial Volume. Am J Psychiatry 176, 228–238, doi:10.1176/appi.ajp.2018.18020149 (2019).

50 Radonjic, N. V. et al. Structural brain imaging studies offer clues about the effects of the shared genetic etiology among neuropsychiatric disorders. Mol Psychiatry 26, 2101–2110, doi:10.1038/s41380-020-01002-z (2021).

51 Castellanos, F. X. et al. Anatomic brain abnormalities in monozygotic twins discordant for attention deficit hyperactivity disorder. American Journal of Psychiatry 160, 1693–1696 (2003).

52 van’t Ent, D., et al. A structural MRI study in monozygotic twins concordant or discordant for attention/hyperactivity problems: Evidence for genetic and environmental heterogeneity in the developing brain. NeuroImage 35, 1004–1020, doi:10.1016/j.neuroimage.2007.01.037 (2007).

53 Saudino, K. J., Ronald, A. & Plomin, R. The etiology of behavior problems in 7-year-old twins: substantial genetic influence and negligible shared environmental influence for parent ratings and ratings by same and different teachers. Journal of abnormal child psychology 33, 113–130 (2005).

54 Hoogman, M. et al. Brain imaging of the cortex in ADHD: a coordinated analysis of large-scale clinical and population-based samples. American Journal of Psychiatry 176, 531–542 (2019).

55 Casey, B. et al. The adolescent brain cognitive development (ABCD) study: imaging acquisition across 21 sites. Developmental cognitive neuroscience 32, 43–54 (2018).

56 Achenbach, T. M. et al. Multicultural assessment of child and adolescent psychopathology with ASEBA and SDQ instruments: research findings, applications, and future directions. Journal of Child Psychology and Psychiatry 49, 251–275 (2008).

57 Chen, W. J., Faraone, S. V., Biederman, J. & Tsuang, M. T. Diagnostic accuracy of the Child Behavior Checklist scales for attention-deficit hyperactivity disorder: a receiver-operating characteristic analysis. Journal of consulting and clinical psychology 62, 1017 (1994).

58 Chang, L.-Y., Wang, M.-Y. & Tsai, P.-S. Diagnostic accuracy of rating scales for attention-deficit/hyperactivity disorder: a meta-analysis. Pediatrics 137, e20152749 (2016).

59 Barch, D. M. et al. Demographic, physical and mental health assessments in the adolescent brain and cognitive development study: Rationale and description. Developmental cognitive neuroscience 32, 55–66 (2018).

60 Achenbach, T. M. DSM-oriented guide for the Achenbach System of Empirically Based Assessment (ASEBA). (ASEBA, 2013).

61 Barch, D. M. et al. Function in the human connectome: task-fMRI and individual differences in behavior. Neuroimage 80, 169–189 (2013).

62 Ashburner, J. A fast diffeomorphic image registration algorithm. Neuroimage 38, 95–113 (2007).

63 Chen, Y. et al. Gray matter volumetric correlates of dimensional impulsivity traits in children: Sex differences and heritability. Human Brain Mapping 43 (2022).

64 Wilke, M., Holland, S. K., Altaye, M. & Gaser, C. Template-O-Matic: a toolbox for creating customized pediatric templates. Neuroimage 41, 903–913, doi:10.1016/j.neuroimage.2008.02.056 (2008).

65 Ide, J. S. et al. Gray matter volumetric correlates of behavioral activation and inhibition system traits in children: An exploratory voxel-based morphometry study of the ABCD project data. Neuroimage 220, 117085, doi:10.1016/j.neuroimage.2020.117085 (2020).

66 Chen, Y. & Li, C.-S. R. Striatal gray matter volumes, externalizing traits, and N-back task performance: An exploratory study of sex differences using the human connectome project data. Journal of Experimental Psychopathology 13, 20438087221080057 (2022).

67 Manjon, J. V., Coupe, P., Marti-Bonmati, L., Collins, D. L. & Robles, M. Adaptive non-local means denoising of MR images with spatially varying noise levels. Journal of magnetic resonance imaging : JMRI 31, 192–203 (2010).

68 Rajapakse, J. C., Giedd, J. N. & Rapoport, J. L. Statistical approach to segmentation of single-channel cerebral MR images. IEEE transactions on medical imaging 16, 176–186 (1997).

69 Gaser, C. & Dahnke, R. CAT-a computational anatomy toolbox for the analysis of structural MRI data. HBM 2016, 336–348 (2016).

70 Cohen, J. Statistical power analysis for the behavioral sciences. (Academic press, 2013).

71 Kleber, B. et al. Voxel-based morphometry in opera singers: Increased gray-matter volume in right somatosensory and auditory cortices. Neuroimage 133, 477–483 (2016).

72 Lakens, D. Calculating and reporting effect sizes to facilitate cumulative science: a practical primer for t-tests and ANOVAs. Frontiers in psychology 4, 863 (2013).

73 Hentschke, H. & Stüttgen, M. C. Computation of measures of effect size for neuroscience data sets. European Journal of Neuroscience 34, 1887–1894 (2011).

74 Zar, J. H. Biostatistical analysis. (Pearson Education India, 1999).

75 Le, T. M. et al. Reward sensitivity and electrodermal responses to actions and outcomes in a go/no-go task. Plos one 14, e0219147 (2019).

76 Dhingra, I. et al. The effects of age on reward magnitude processing in the monetary incentive delay task. NeuroImage 207, 116368 (2020).

77 Ide, J. S. et al. Gray matter volumetric correlates of behavioral activation and inhibition system traits in children: An exploratory voxel-based morphometry study of the ABCD project data. Neuroimage 220, 117085 (2020).

78 Chen, Y., Chaudhary, S., Wang, W. & Li, C.-S. R. Gray matter volumes of the insula and anterior cingulate cortex and their dysfunctional roles in cigarette smoking. Addiction Neuroscience, 100003 (2021).

79 Li, G. et al. Sex Differences in Neural Responses to the Perception of Social Interactions. Front Hum Neurosci 14, 565132, doi:10.3389/fnhum.2020.565132 (2020).

80 Visscher, P. M. et al. Assumption-free estimation of heritability from genome-wide identity-by-descent sharing between full siblings. PLoS Genet 2, e41 (2006).

81 Visscher, P. M., Hill, W. G. & Wray, N. R. Heritability in the genomics era— concepts and misconceptions. Nature reviews genetics 9, 255–266 (2008).

82 Muthen, L. K. & Muthen, B. O. Mplus user’s guide. 7th. Los Angeles, CA: Muthén & Muthén 19982006 (2012).

83 Kohler, H.-P., Behrman, J. R. & Schnittker, J. Social science methods for twins data: Integrating causality, endowments, and heritability. Biodemography and Social Biology 57, 88–141 (2011).

84 Rijsdijk, F. V. & Sham, P. C. Analytic approaches to twin data using structural equation models. Briefings in bioinformatics 3, 119–133 (2002).

85 Chen, Y. et al. Accessing the development and heritability of the capacity of cognitive control. Neuropsychologia 139, 107361 (2020).

86 Baroncini, M. et al. MRI atlas of the human hypothalamus. Neuroimage 59, 168–180, doi:10.1016/j.neuroimage.2011.07.013 (2012).

87 Mai, J. K., Paxinos, G. & Voss, T. Atlas of the human brain. 3rd edn, (Academic Press, 2008).

88 Satizabal, C. L. et al. Genetic architecture of subcortical brain structures in 38,851 individuals. Nat Genet 51, 1624–1636, doi:10.1038/s41588-019-0511-y (2019).

89 Hesslinger, B. et al. Frontoorbital volume reductions in adult patients with attention deficit hyperactivity disorder. Neuroscience letters 328, 319–321 (2002).

90 Krain, A. L. & Castellanos, F. X. Brain development and ADHD. Clinical psychology review 26, 433–444 (2006).

91 Fernández-Jaén, A. et al. Cortical thinning of temporal pole and orbitofrontal cortex in medication-naive children and adolescents with ADHD. Psychiatry Research: Neuroimaging 224, 8–13 (2014).

92 Rubia, K. “Cool” inferior frontostriatal dysfunction in attention-deficit/hyperactivity disorder versus “hot” ventromedial orbitofrontal-limbic dysfunction in conduct disorder: a review. Biological psychiatry 69, e69–e87 (2011).

93 Fan, J. et al. Quantitative characterization of functional anatomical contributions to cognitive control under uncertainty. Journal of cognitive neuroscience 26, 1490–1506 (2014).

94 Wu, T. et al. Hick–Hyman law is mediated by the cognitive control network in the brain. Cerebral Cortex 28, 2267–2282 (2018).

95 Kuntsi, J., Rijsdijk, F., Ronald, A., Asherson, P. & Plomin, R. Genetic influences on the stability of attention-deficit/hyperactivity disorder symptoms from early to middle childhood. Biological Psychiatry 57, 647–654 (2005).

96 Fuermaier, A. et al. Cognitive impairment in adult ADHD—perspective matters! Neuropsychology 29, 45 (2015).

97 Fairchild, G. Hypothalamic-pituitary-adrenocortical axis function in attention-deficit hyperactivity disorder. Current topics in behavioral neurosciences 9, 93–111, doi:10.1007/7854_2010_101 (2012).

98 Chang, J. P. et al. Cortisol, inflammatory biomarkers and neurotrophins in children and adolescents with attention deficit hyperactivity disorder (ADHD) in Taiwan. Brain Behav Immun 88, 105–113, doi:10.1016/j.bbi.2020.05.017 (2020).

99 Angeli, E. et al. Salivary cortisol and alpha-amylase diurnal profiles and stress reactivity in children with Attention Deficit Hyperactivity Disorder. Psychoneuroendocrinology 90, 174–181, doi:10.1016/j.psyneuen.2018.02.026 (2018).

100 Korpa, T. et al. Mothers’ parenting stress is associated with salivary cortisol profiles in children with attention deficit hyperactivity disorder. Stress 20, 149–158, doi:10.1080/10253890.2017.1303472 (2017).

101 Chevrier, A. & Schachar, R. J. BOLD differences normally attributed to inhibitory control predict symptoms, not task-directed inhibitory control in ADHD. J Neurodev Disord 12, 8, doi:10.1186/s11689-020-09311-8 (2020).

102 Modi, S., Thaploo, D., Kumar, P. & Khushu, S. Individual differences in trait anxiety are associated with gray matter alterations in hypothalamus: Preliminary neuroanatomical evidence. Psychiatry Res Neuroimaging 283, 45–54, doi:10.1016/j.pscychresns.2018.11.008 (2019).

103 Carmona, S. et al. Global and regional gray matter reductions in ADHD: A voxel-based morphometric study. Neuroscience Letters 389, 88–93, doi:10.1016/j.neulet.2005.07.020 (2005).

104 Wu, Z.-M. et al. Linked anatomical and functional brain alterations in children with attention-deficit/hyperactivity disorder. NeuroImage: Clinical 23, 101851 (2019).

105 Zhang-James, Y. et al. Evidence for similar structural brain anomalies in youth and adult attention-deficit/hyperactivity disorder: a machine learning analysis. Translational psychiatry 11, 1–9 (2021).

106 Shang, C., Lin, H., Tseng, W. & Gau, S. A haplotype of the dopamine transporter gene modulates regional homogeneity, gray matter volume, and visual memory in children with attention-deficit/hyperactivity disorder. Psychological medicine 48, 2530–2540 (2018).

107 Wang, L.-J. et al. Gray matter volume and microRNA levels in patients with attention-deficit/hyperactivity disorder. European archives of psychiatry and clinical neuroscience 270, 1037–1045 (2020).

108 Shen, C. et al. Neural correlates of the dual-pathway model for ADHD in adolescents. American Journal of Psychiatry 177, 844–854 (2020).

109 Chang, J.-C., Lin, H.-Y., Lv, J., Tseng, W.-Y. I. & Gau, S. S.-F. Regional brain volume predicts response to methylphenidate treatment in individuals with ADHD. BMC psychiatry 21, 1–14 (2021).

110 Ide, J. S., Tung, H. C., Yang, C.-T., Tseng, Y.-C. & Li, C.-S. R. Barratt Impulsivity in Healthy Adults Is Associated with Higher Gray Matter Concentration in the Parietal Occipital Cortex that Represents Peripheral Visual Field. Frontiers in Human Neuroscience 11, doi:10.3389/fnhum.2017.00222 (2017).

111 Castellanos, F. X. & Proal, E. Large-scale brain systems in ADHD: beyond the prefrontal–striatal model. Trends in Cognitive Sciences 16, 17–26, doi:10.1016/j.tics.2011.11.007 (2012).

112 Dirlikov, B. et al. Distinct frontal lobe morphology in girls and boys with ADHD. Neuroimage: Clinical 7, 222–229 (2015).

113 Fan, J. An information theory account of cognitive control. Frontiers in human neuroscience 8, 680 (2014).

114 Wu, T. et al. Anterior insular cortex is a bottleneck of cognitive control. NeuroImage 195, 490–504 (2019).

115 Gaub, M. & Carlson, C. L. Gender differences in ADHD: a meta-analysis and critical review. Journal of the American Academy of Child & Adolescent Psychiatry 36, 1036–1045 (1997).

116 Gershon, J. & Gershon, J. A meta-analytic review of gender differences in ADHD. Journal of attention disorders 5, 143–154 (2002).

117 Gross-Tsur, V., et al. The impact of sex and subtypes on cognitive and psychosocial aspects of ADHD. Developmental Medicine & Child Neurology 48, 901–905 (2006).

118 Li, T. et al. Characterizing neuroanatomic heterogeneity in people with and without ADHD based on subcortical brain volumes. Journal of Child Psychology and Psychiatry (2021).

119 Hicks, B. M. et al. Gender differences and developmental change in externalizing disorders from late adolescence to early adulthood: A longitudinal twin study. Journal of abnormal psychology 116, 433 (2007).

120 Zucker. Anticipating problem alcohol use developmentally from childhood into middle adulthood: what have we learned? Addiction 103, 100–108 (2008).

121 Chen, Y., Li, G., Ide, J. S., Luo, X. & Li, C.-S. R. Sex differences in attention deficit hyperactivity symptom severity and functional connectivity of the dorsal striatum in young adults. Neuroimage: Reports 1, doi:10.1016/j.ynirp.2021.100025 (2021).

122 Sobel, L. J. et al. Basal ganglia surface morphology and the effects of stimulant medications in youth with attention deficit hyperactivity disorder. American Journal of Psychiatry 167, 977–986 (2010).

123 Herskovits, E. H. et al. Is the spatial distribution of brain lesions associated with closed-head injury predictive of subsequent development of attention-deficit/hyperactivity disorder? Analysis with brain-image database. Radiology 213, 389–394 (1999).

124 Max, J. E. et al. Putamen lesions and the development of attention-deficit/hyperactivity symptomatology. Journal of the American Academy of Child & Adolescent Psychiatry 41, 563–571 (2002).

125 Faraone, S. V. & Biederman, J. Neurobiology of attention-deficit hyperactivity disorder. Biological psychiatry 44, 951–958 (1998).

126 Tuvblad, C., Zheng, M., Raine, A. & Baker, L. A. A common genetic factor explains the covariation among ADHD ODD and CD symptoms in 9–10 year old boys and girls. Journal of abnormal child psychology 37, 153–167 (2009).

127 Wood, A. C. & Neale, M. C. Twin studies and their implications for molecular genetic studies: endophenotypes integrate quantitative and molecular genetics in ADHD research. Journal of the American Academy of Child & Adolescent Psychiatry 49, 874–883 (2010).

128 Wolf, L. E. & Wasserstein, J. Adult ADHD: concluding thoughts. ANNALS-NEW YORK ACADEMY OF SCIENCES 931, 396–408 (2001).

129 Brassett-Harknett, A. & Butler, N. Attention-deficit/hyperactivity disorder: An overview of the etiology and a review of the literature relating to the correlates and lifecourse outcomes for men and women. Clinical Psychology Review 27, 188–210, doi:10.1016/j.cpr.2005.06.001 (2007).

130 Peterson, B. S., Pine, D. S., Cohen, P. & Brook, J. S. Prospective, longitudinal study of tic, obsessive-compulsive, and attention-deficit/hyperactivity disorders in an epidemiological sample. Journal of the American Academy of Child & Adolescent Psychiatry 40, 685–695 (2001).

131 Hudziak, J. J., Derks, E. M., Althoff, R. R., Rettew, D. C. & Boomsma, D. I. The genetic and environmental contributions to attention deficit hyperactivity disorder as measured by the Conners’ Rating Scales—Revised. American Journal of Psychiatry 162, 1614–1620 (2005).

132 Swagerman, S. C., Brouwer, R. M., de Geus, E. J., Hulshoff Pol, H. E. & Boomsma, D. I. Development and heritability of subcortical brain volumes at ages 9 and 12. Genes Brain Behav 13, 733–742, doi:10.1111/gbb.12182 (2014).

133 Wallace, G. L. et al. A bivariate twin study of regional brain volumes and verbal and nonverbal intellectual skills during childhood and adolescence. Behavior genetics 40, 125–134 (2010).

134 Yoon, U., Perusse, D., Lee, J.-M. & Evans, A. C. Genetic and environmental influences on structural variability of the brain in pediatric twin: deformation based morphometry. Neuroscience letters 493, 8–13 (2011).

135 Stein, J. L. et al. Discovery and replication of dopamine-related gene effects on caudate volume in young and elderly populations (N=1198) using genome-wide search. Mol Psychiatry 16, 927–937, 881, doi:10.1038/mp.2011.32 (2011).

136 Kremen, W. S. et al. Genetic and environmental influences on the size of specific brain regions in midlife: the VETSA MRI study. Neuroimage 49, 1213–1223 (2010).

137 den Braber, A., et al. Heritability of subcortical brain measures: a perspective for future genome-wide association studies. Neuroimage 83, 98–102, doi:10.1016/j.neuroimage.2013.06.027 (2013).

138 Blokland, G. A. et al. Quantifying the heritability of task-related brain activation and performance during the N-back working memory task: a twin fMRI study. Biol Psychol 79, 70–79, doi:10.1016/j.biopsycho.2008.03.006 (2008).

139 Briley, D. A. & Tucker-Drob, E. M. Explaining the increasing heritability of cognitive ability across development: a meta-analysis of longitudinal twin and adoption studies. Psychol Sci 24, 1704–1713, doi:10.1177/0956797613478618 (2013).

140 Franke, B. et al. The genetics of attention deficit/hyperactivity disorder in adults, a review. Molecular psychiatry 17, 960–987 (2012).

141 Ehringer, M. A., Rhee, S. H., Young, S., Corley, R. & Hewitt, J. K. Genetic and environmental contributions to common psychopathologies of childhood and adolescence: a study of twins and their siblings. J Abnorm Child Psychol 34, 1–17, doi:10.1007/s10802-005-9000-0 (2006).

142 Friedman, L. A. & Rapoport, J. L. Brain development in ADHD. Current opinion in neurobiology 30, 106–111 (2015).

143 Daviss, W. B. A review of co-morbid depression in pediatric ADHD: etiologies, phenomenology, and treatment. Journal of child and adolescent psychopharmacology 18, 565–571 (2008).

144 Rogers, J. C. & De Brito, S. A. Cortical and subcortical gray matter volume in youths with conduct problems: a meta-analysis. JAMA psychiatry 73, 64–72 (2016).

145 Boedhoe, P. S. et al. Subcortical brain volume, regional cortical thickness, and cortical surface area across disorders: Findings from the ENIGMA ADHD, ASD, and OCD working groups. American Journal of Psychiatry 177, 834–843 (2020).

146 Hoogman, M. et al. Consortium neuroscience of attention deficit/hyperactivity disorder and autism spectrum disorder: The ENIGMA adventure. Human brain mapping (2020).

147 Opel, N. et al. Cross-disorder analysis of brain structural abnormalities in six major psychiatric disorders: a secondary analysis of mega-and meta-analytical findings from the ENIGMA consortium. Biological Psychiatry 88, 678–686 (2020).

148 Thijssen, S., Collins, P. F. & Luciana, M. Pubertal development mediates the association between family environment and brain structure and function in childhood. Dev Psychopathol 32, 687–702, doi:10.1017/S0954579419000580 (2020).

149 Cheng, T. W. et al. A Researcher’s Guide to the Measurement and Modeling of Puberty in the ABCD Study((R)) at Baseline. Front Endocrinol (Lausanne) 12, 608575, doi:10.3389/fendo.2021.608575 (2021).

150 Assary, E., Zavos, H. M. S., Krapohl, E., Keers, R. & Pluess, M. Genetic architecture of Environmental Sensitivity reflects multiple heritable components: a twin study with adolescents. Mol Psychiatry 26, 4896–4904, doi:10.1038/s41380-020-0783-8 (2021).

