## Supplement for "Gray Matter Volumetric Correlates of Attention Deficit and Hyperactivity Traits in Emerging Adolescents"

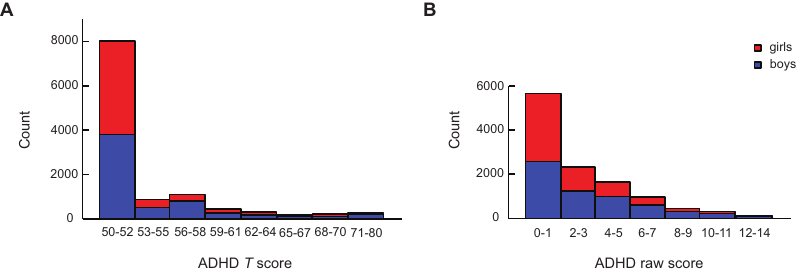

**Supplementary Figure S1.** Distributions of (**A**) ADHD *T* score and (**B**) ADHD raw score of boys and girls.

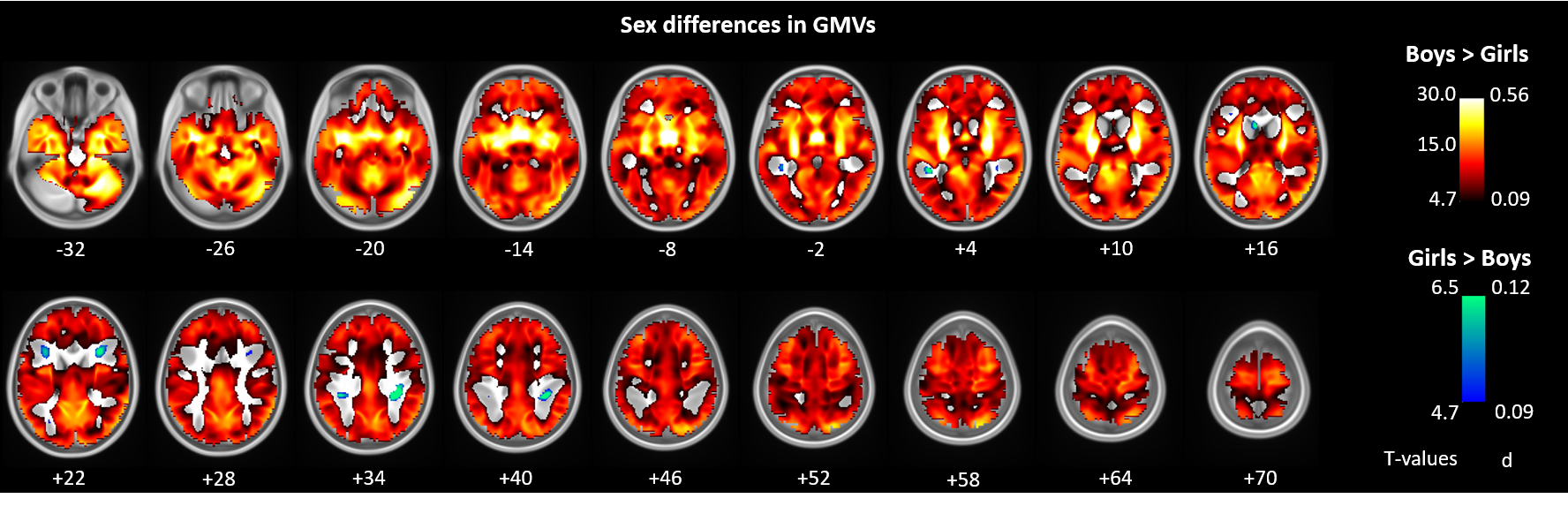

**Supplementary Figure S2.** Sex differences in GMVs. Two-sample *t* test, voxel *p* < 0.05, FWE corrected. Color bars show the voxel *T* and the corresponding effect size (*d*) values.

**Supplementary Table S1.** GMV correlates of ADHD scores for girls and boys combined and separately. Whole-brain regression, *p* < 0.05, FWE corrected.

| **Cluster size (k)** | **Voxel Z-score** | **MNI Coordinates (mm)** | | | **Identified Region** |
| --- | --- | --- | --- | --- | --- |
|  |  | **x** | **y** | **z** |  |
| ***All*** |  |  |  |  |  |
| 689 | 6.88 | -5 | -13 | -9 | Hypothalamus |
| 48608 | -7.63 | -13 | 2 | 6 | Frontal_Mid, Frontal_Mid_Orb, Caudate, Putamen (L+R); Insula_L |
| 19723 | -7.59 | -32 | -99 | 6 | Occipital_Mid_L |
| 7543 | -7.34 | 16 | 10 | 4 | Insula_R, Putamen_R |
| 3071 | -6.67 | -27 | -21 | -20 | ParaHipp_L, Hippocampus_L, Fusiform_L |
| 3931 | -6.63 | -25 | -67 | -13 | Lingual_L, Cerebellum_L |
| 4298 | -6.37 | -68 | -32 | 29 | Parietal_Inf_L |
| 6276 | -6.33 | 43 | -92 | -2 | Occipital_Mid_R |
| 1837 | -6.25 | 23 | -9 | -23 | ParaHippocampal_R |
| 143 | -6.19 | 21 | 23 | -23 | Frontal_Inf_Orb_R |
| 2929 | -6.19 | 70 | -24 | 24 | SupraMarginal_R |
| 4146 | -6.11 | 70 | -11 | -23 | Temporal_Inf_R |
| 3245 | -6.07 | -38 | -24 | 37 | Postcentral_L |
| 3681 | -6.02 | 60 | -6 | 40 | Postcentral_R, Frontal_Mid_R |
| 575 | -5.82 | 22 | -37 | -12 | ParaHippocampal_R |
| 1424 | -5.71 | 62 | 14 | 33 | Precentral_R, Frontal_Inf_Oper_R |
| 3762 | -5.6 | -4 | -66 | 16 | Calcarine_L, Cuneus_L |
| 1575 | -5.58 | -24 | 26 | 61 | Frontal_Sup_L, Frontal_Sup_Medial_L |
| 790 | -5.52 | 29 | -59 | -12 | Fusiform_R |
| 1286 | -5.47 | -52 | -1 | 25 | Precentral_L |
| 2849 | -5.44 | 44 | 5 | -37 | Temporal_Inf_R |
| 407 | -5.4 | -39 | -51 | 50 | Parietal_Inf_L |
| 1936 | -5.32 | -62 | -9 | -15 | Temporal_Mid_L, Temporal_Sup_L |
| 1339 | -5.31 | 11 | 44 | -5 | Frontal_Med_Orb_R |
| 469 | -5.23 | -38 | 12 | -23 | Temporal_Pole_Sup_L |
| 299 | -5.19 | -54 | 30 | 14 | Frontal_Inf_Tri_L |
| 291 | -5.16 | -12 | -50 | 61 | Precuneus_L |
| 392 | -5.15 | -8 | -62 | 56 | Precuneus_L |
| 328 | -5.15 | 9 | -71 | 41 | Precuneus_R |
| 171 | -5.07 | 10 | -18 | 35 | Cingulum_Mid_R |
| 335 | -5.04 | -11 | 50 | 47 | Frontal_Sup_Medial_L |
| 627 | -5.03 | -6 | 36 | 21 | ACC |
| 104 | -4.92 | -39 | 41 | 35 | Frontal_Mid_L |
| 225 | -4.9 | 61 | -60 | -10 | Temporal_Inf_R |
| 37 | -4.9 | 46 | -52 | 18 | Temporal_Mid_R |
| 130 | -4.85 | 59 | -62 | 34 | Angular_R |
| 116 | -4.85 | 54 | 12 | 6 | Frontal_Inf_Oper_R |
| 204 | -4.85 | 26 | -60 | -26 | Cerebelum_6_R |
| 135 | -4.77 | 43 | -50 | 51 | Parietal_Inf_R |
| 216 | -4.76 | -47 | -24 | -22 | Temporal_Inf_L |
| 32 | -4.74 | 19 | -53 | 11 | Calcarine_R |
| 158 | -4.73 | 4 | -13 | 46 | Cingulum_Mid_R |
| 122 | -4.72 | 40 | -22 | -25 | Fusiform_R |
| 62 | -4.72 | 44 | -34 | 42 | SupraMarginal_R |
| 37 | -4.59 | 68 | -12 | 10 | Temporal_Sup_R |
| ***Girls*** |  |  |  |  |  |
| 19966 | -6.36 | -26 | -97 | 7 | Occipital_Mid_L, Occipital_Sup_L |
| 10056 | -6.17 | 40 | 8 | -45 | Temporal_Inf_R, Temporal_Pole_Mid_R |
| 10449 | -6.02 | -34 | 4 | -44 | Temporal_Pole_Mid_L, Temporal_Inf_L |
| 3697 | -5.89 | -25 | -69 | -15 | Fusiform_L, Cerebelum_L |
| 2325 | -5.73 | -27 | -19 | -21 | Hippocampus_L, ParaHippocampal_L |
| 1487 | -5.65 | -23 | 19 | -1 | Putamen_L |
| 1431 | -5.52 | 10 | 14 | -3 | Caudate_R |
| 2516 | -5.47 | 61 | -62 | 36 | Temporal_Sup_R, |
| 1360 | -5.41 | 30 | -18 | -22 | ParaHippocampal_R |
| 1005 | -5.38 | 70 | -30 | 28 | SupraMarginal_R |
| 1550 | -5.3 | 16 | -75 | 62 | Parietal_Sup_R |
| 431 | -5.1 | -33 | 59 | 16 | Frontal_Mid_L |
| 1434 | -5.1 | 71 | -15 | -21 | Temporal_Mid_R |
| 613 | -5.1 | 46 | 12 | -6 | Insula_R, Frontal_Inf_Oper_R |
| 1354 | -5.1 | 48 | -72 | 22 | Temporal_Mid_R |
| 785 | -5.09 | -9 | 9 | -3 | Caudate_L |
| 300 | -5.08 | -26 | -65 | 67 | Parietal_Sup_L |
| 409 | -5.05 | 61 | -59 | -8 | Temporal_Inf_R |
| 390 | -4.96 | -54 | -64 | 19 | Temporal_Mid_L |
| 277 | -4.92 | -66 | -54 | 26 | SupraMarginal_L |
| 600 | -4.91 | -42 | 50 | -12 | Frontal_Inf_Orb_L, Frontal_Mid_Orb_L |
| 133 | -4.88 | 46 | -68 | -16 | Occipital_Inf_R |
| 497 | -4.86 | 39 | 61 | 5 | Frontal_Mid_R |
| 62 | -4.84 | 52 | 12 | 15 | Frontal_Inf_Oper_R |
| 190 | -4.83 | 22 | -59 | 12 | Calcarine_R |
| 38 | -4.8 | 20 | -86 | 48 | Cuneus_R |
| 64 | -4.79 | 7 | 60 | -21 | Frontal_Sup_Orb_R |
| 39 | -4.75 | 47 | -52 | 16 | Temporal_Mid_R |
| 73 | -4.72 | 20 | -102 | -2 | Calcarine_R |
| 73 | -4.68 | 68 | -36 | 3 | Temporal_Mid_R |
| 39 | -4.66 | 10 | -61 | 2 | Lingual_R |
| 191 | -4.66 | -68 | -22 | 31 | SupraMarginal_L, Postcentral_L |
| 63 | -4.66 | -38 | 42 | 37 | Frontal_Mid_L |
| 91 | -4.64 | -51 | -2 | 26 | Precentral_L |
| 39 | -4.6 | -44 | -71 | 29 | Occipital_Mid_L |
| ***Boys*** |  |  |  |  |  |
| 103945 | -7.3 | -23 | 18 | 3 | Putamen_L, Frontal_Mid_Orb_R |
| 15808 | -6.85 | -34 | -98 | 7 | Occipital_Mid_L |
| 9291 | -6.42 | 44 | -91 | -1 | Occipital_Mid_R |
| 3906 | -5.94 | 60 | -6 | 42 | Postcentral_R, Frontal_Inf_Oper_R |
| 1682 | -5.92 | -27 | -65 | -31 | Cerebelum_6_L |
| 3477 | -5.91 | -17 | 22 | 66 | Frontal_Sup_L |
| 8443 | -5.87 | 70 | -11 | -25 | Temporal_Inf_R |
| 9858 | -5.67 | -50 | -26 | 15 | Rolandic_Oper_L, Temporal_Mid_L |
| 2149 | -5.62 | -4 | -65 | 15 | Calcarine_L |
| 2454 | -5.59 | -29 | -56 | -50 | Cerebelum_8_L |
| 676 | -5.5 | 20 | -78 | -50 | Cerebelum_7b_R |
| 1862 | -5.49 | 62 | -22 | 22 | SupraMarginal_R |
| 1664 | -5.43 | 43 | -73 | 49 | Angular_R, Parietal_Sup_R |
| 905 | -5.43 | 27 | -64 | -29 | Cerebelum_6_R |
| 1431 | -5.39 | -52 | -6 | 50 | Precentral_L |
| 277 | -5.38 | -52 | -14 | -2 | Temporal_Sup_L |
| 669 | -5.37 | -6 | 34 | 23 | ACC |
| 326 | -5.37 | -8 | -23 | 32 | Cingulum_Post_L |
| 344 | -5.28 | -10 | 35 | 41 | Frontal_Sup_Medial_L |
| 379 | -5.27 | 34 | 23 | 60 | Frontal_Mid_R |
| 311 | -5.25 | 24 | -38 | -10 | ParaHippocampal_R |
| 105 | -5.14 | -56 | -67 | 24 | Temporal_Mid_L |
| 315 | -5.08 | 27 | -59 | -12 | Fusiform_R |
| 243 | -5.07 | 10 | -65 | 43 | Precuneus_R |
| 307 | -5.04 | -64 | -50 | 32 | SupraMarginal_L, Temporal_Sup_L |
| 344 | -4.93 | -24 | -72 | -14 | Fusiform_L, Lingual_L |
| 193 | -4.91 | 39 | -28 | -18 | Fusiform_R |
| 121 | -4.89 | -48 | -23 | -28 | Temporal_Inf_L |
| 224 | -4.87 | -5 | -64 | 53 | Precuneus_L |
| 54 | -4.87 | 37 | -57 | -48 | Cerebelum_8_R |
| 143 | -4.83 | 29 | -85 | -41 | Cerebelum_Crus2_R |
| 85 | -4.82 | 17 | 25 | 65 | Frontal_Sup_R |
| 98 | -4.79 | -52 | -65 | -15 | Occipital_Inf_L |
| 97 | -4.76 | -5 | -58 | -22 | Vermis_6 |
| 93 | -4.75 | -64 | -59 | -18 | Temporal_Inf_L |
| 126 | -4.75 | 1 | -26 | 45 | Cingulum_Mid_R |
| 59 | -4.7 | 10 | -47 | 52 | Precuneus_R |
| 37 | -4.7 | -51 | -48 | 27 | SupraMarginal_L |
| 77 | -4.66 | -62 | -25 | -25 | Temporal_Inf_L |
| 195 | -4.66 | 39 | -3 | 5 | Insula_R |
| 41 | -4.65 | -14 | -73 | 64 | Precuneus_L |

*Note*: Brain regions are identified with the Automated Anatomic Labeling atlas (Tzourio-Mazoyer et al., 2002) except for clusters*, which are identified by consulting an atlas (Duvernoy, 2003). The sign of Z value indicates the direction of correlation. R: right; L: left; sup: superior; mid: middle; inf: inferior; G: gyrus

**Supplementary Table S2.** Slope tests of sex differences in the correlation of GMVs vs. 2-back accuracy rates.

| **ROI** | **Slope-T** | **p-value** | **Correlation for girls** | **Correlations for boys** |
| --- | --- | --- | --- | --- |
| Hypothalamus | 0.211 | 0.833 | *r* = 0.035, *p* = 0.013 | *r* = 0.039, *p* = 0.005 |
| Girls mask | 0.543 | 0.587 | *r* = 0.190, *p* < 0.001 | *r* = 0.173, *p* < 0.001 |
| Boys mask | 0.274 | 0.784 | *r* = 0.154, *p* < 0.001 | *r* = 0.151, *p* < 0.001 |
| Girls spec. mask | 0.612 | 0.541 | *r* = 0.189, *p* < 0.001 | *r* = 0.172, *p* < 0.001 |
| Boys spec. mask | 0.351 | 0.726 | *r* = 0.150, *p* < 0.001 | *r* = 0.148, *p* < 0.001 |
| Caudate (bilat) | 1.858 | 0.063 | *r* = 0.113, *p* < 0.001 | *r* = 0.072, *p* < 0.001 |
| Putamen (bilat) | 0.832 | 0.406 | *r* = 0.100, *p* < 0.001 | *r* = 0.082, *p* < 0.001 |
| Pallidum (bilat) | 0.781 | 0.435 | *r* = 0.085, *p* < 0.001 | *r* = 0.066, *p* < 0.001 |

*Note:* **p* < 0.05/8 = 0.00625; spec.: specific; bilat: bilateral; Age, race, study site, twin status, TIV, and scanner model were included as covariates.

**Supplementary Table S3.** Slope tests of sex differences in the correlation of GMVs vs. ADHD scores.

| **ROI** | **Slope-T** | **p-value** | **Correlation for girls** | **Correlations for boys** |
| --- | --- | --- | --- | --- |
| Hypothalamus | 0.250 | 0.803 | *r* = 0.024, *p* = 0.077 | *r* = 0.026, *p* = 0.044 |
| Girls mask | 0.650 | 0.516 | *r* = -0.108, *p* < 0.001 | *r* = -0.085, *p* < 0.001 |
| Boys mask | 2.788 | 0.005* | *r* = -0.072, *p* < 0.001 | *r* = -0.112, *p* < 0.001 |
| Girls spec. mask | 1.156 | 0.248 | *r* = -0.107, *p* < 0.001 | *r* = -0.076, *p* < 0.001 |
| Boys spec. mask | 2.990 | 0.003* | *r* = -0.068, *p* < 0.001 | *r* = -0.112, *p* < 0.001 |
| Caudate (bilat) | 0.406 | 0.685 | *r* = -0.056, *p* < 0.001 | *r* = -0.057, *p* < 0.001 |
| Putamen (bilat) | 1.064 | 0.288 | *r* = -0.045, *p* = 0.001 | *r* = -0.061, *p* < 0.001 |
| Pallidum (bilat) | 1.408 | 0.159 | *r* = -0.052, *p* < 0.001 | *r* = -0.071, *p* < 0.001 |

*Note:* **p* < 0.05/8 = 0.00625; excl: exclusive; bilat: bilateral; Age, race, study site, twin/sibling status, TIV, and scanner model were included as covariates.

**Supplementary Table S4.** Linear mixed-effects models of GMV correlates with an interaction term of sex and ADHD score.

| **ROI** | *Sex* | | *ADHD* | | *Sex × ADHD* | |
| --- | --- | --- | --- | --- | --- | --- |
|  | *β (SE)* | *p* | *β (SE)* | *p* | *β (SE)* | *p* |
| Hypothalamus | -0.017 (0.007) | 0.008 | 0.002 (0.0001) | 0.005 | 0.000009 (0.0001) | 0.941 |
| Girls mask | -0.010 (0.007) | 0.142 | -0.001  (0.0001) | < 0.001 | -0.0002 (0.0001) | 0.172 |
| Boys mask | -0.026 (0.006) | < 0.001 | -0.001 (0.0001) | < 0.001 | 0.0002 (0.0001) | 0.057 |
| Girls spec. mask | -0.008 (0.007) | 0.279 | -0.001 (0.0001) | < 0.001 | -0.0002 (0.0001) | 0.076 |
| Boys spec. mask | -0.027 (0.006) | < 0.001 | -0.001 (0.0001) | < 0.001 | 0.0002 (0.0001) | 0.034 |
| Caudate (bilat) | -0.005 (0.007) | 0.469 | -0.0004 (0.0001) | < 0.001 | 0.000000 (0.0001) | 0.998 |
| Putamen (bilat) | -0.026 (0.007) | < 0.001 | -0.0004 (0.0001) | < 0.001 | 0.0001 (0.0001) | 0.362 |
| Pallidum (bilat) | -0.027 (0.009) | 0.005 | -0.0006 (0.0001) | < 0.001 | 0.0002 (0.0002) | 0.325 |

*Note:* The models were estimated with twin/sibling status as a random effect after controlling for age, race, study site, scanner model, and TIV.

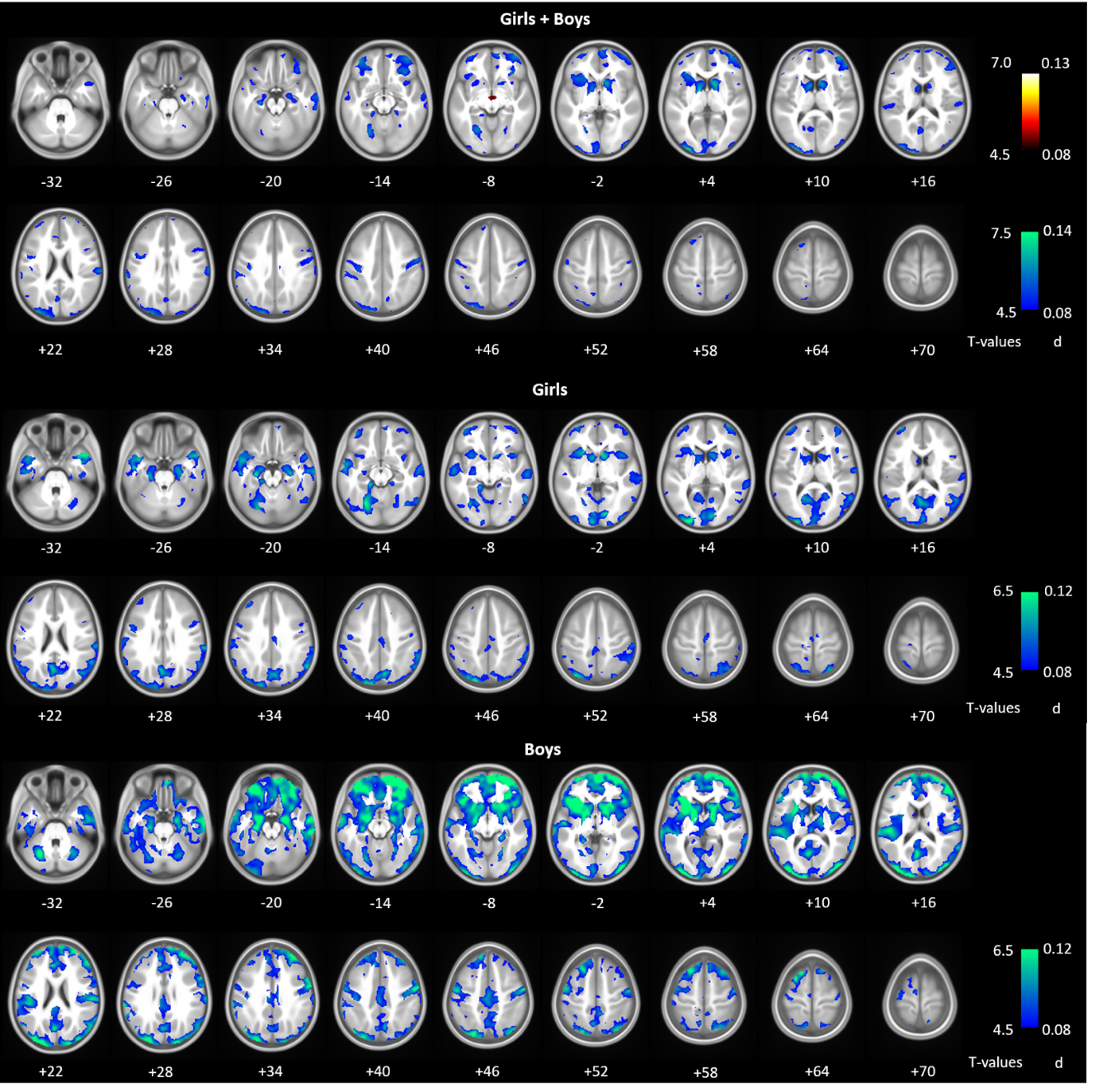

**Supplementary Figure S3.** GMV correlates of ADHD scores for girls and boys combined, and for girls and boys, separately, without TIV included as a covariate. Cool color bars show voxel *T* (negative correlations) and *d* values.

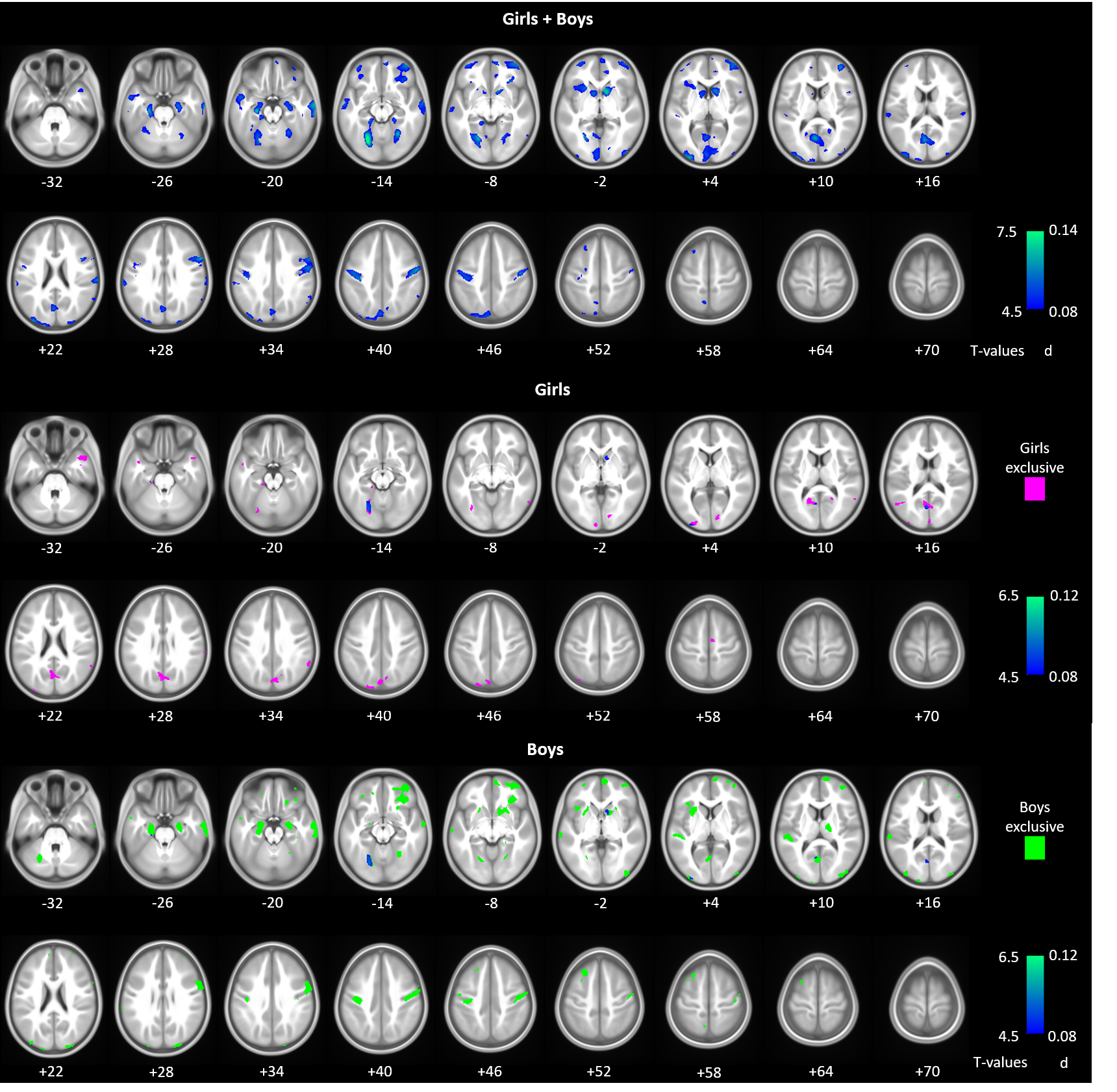

**Supplementary Figure S4.** GMV correlates of ADHD scores (non-medicated children only) for girls and boys combined, and for girls and boys, separately. Cool color bars show voxel *T* (negative correlations) and d values. For girls and boys separately, we performed an exclusive masking to highlight clusters that are specific to girls (pink) and to boys (light green).

**Supplementary Table S5.** GMV correlates of ADHD scores for girls and boys combined and separately (non-medicated children only). Whole-brain regression, *p* < 0.05, FWE corrected.

| **Cluster size (k)** | **Voxel Z-score** | **MNI Coordinates (mm)** | | | **Identified Region** |
| --- | --- | --- | --- | --- | --- |
|  |  | **x** | **y** | **z** |  |
| ***All*** |  |  |  |  |  |
| 40219 | 7.06 | -23 | -70 | -14 | Fusiform_L, ParaHipp_L , Calcarine_L |
| 3162 | 6.59 | 12 | 12 | -3 | Caudate_R |
| 5100 | 6.41 | 28 | -59 | -14 | Fusiform_R, Lingual_L, ParaHipp_L |
| 3606 | 6.41 | 70 | -13 | -23 | Temporal_Inf_R, Temporal_Mid_R |
| 8523 | 6.4 | 62 | 14 | 30 | Frontal_Inf_Oper_R, Postcentral_R |
| 9807 | 6.25 | 39 | 52 | 9 | Frontal_Mid_R, Frontal_Inf_Orb_R |
| 3460 | 6.23 | -23 | 19 | 2 | Putamen_L, Insula_L |
| 1857 | 5.98 | 25 | -12 | -24 | ParaHipp_R, Cerebelum_4_5_R |
| 5249 | 5.94 | -37 | -24 | 41 | Postcentral_L |
| 2163 | 5.73 | -11 | 8 | -1 | Caudate_L |
| 1981 | 5.58 | 11 | 60 | -10 | Frontal_Med_Orb_R, Rectus_R |
| 4646 | 5.58 | -54 | 2 | -22 | Temporal_Mid_L |
| 2276 | 5.57 | 26 | -95 | 29 | Occipital_Mid_R |
| 1572 | 5.55 | -41 | 6 | -44 | Temporal_Pole_Mid_L, Temporal_Inf_L |
| 2503 | 5.5 | -68 | -31 | 27 | Temporal_Sup_L, SupraMarginal_L |
| 748 | 5.46 | -50 | -3 | 26 | Precentral_L |
| 4251 | 5.42 | -39 | 56 | -10 | Frontal_Mid_Orb_L, Frontal_Sup_Orb |
| 1358 | 5.37 | 57 | -60 | 36 | Angular_R |
| 941 | 5.37 | -26 | 26 | 58 | Frontal_Mid_L |
| 644 | 5.32 | -7 | -61 | 57 | Precuneus_L |
| 2840 | 5.23 | 45 | 4 | -45 | Temporal_Inf_R, Temporal_Pole_Mid |
| 870 | 5.21 | 12 | 44 | -4 | Frontal_Med_Orb_R |
| 153 | 5.1 | -11 | 52 | 10 | Frontal_Sup_Medial_L |
| 275 | 5.07 | 21 | -53 | 11 | Calcarine_R |
| 91 | 5.06 | -47 | -24 | -22 | Temporal_Inf_L |
| 415 | 5.03 | 61 | -60 | -10 | Temporal_Inf_R |
| 279 | 4.85 | 13 | 31 | 14 | Cingulum_Ant_R |
| 131 | 4.84 | -52 | -66 | 19 | Temporal_Mid_L |
| 221 | 4.74 | 40 | -25 | -23 | Fusiform_R |
| 235 | 4.73 | 7 | -11 | 61 | Supp_Motor_Area_R |
| 151 | 4.73 | -59 | -22 | -26 | Temporal_Inf_L |
| 102 | 4.73 | -6 | 37 | 19 | Cingulum_Ant_L |
| 34 | 4.66 | -11 | -51 | 60 | Precuneus_L |
| 33 | 4.61 | -50 | -75 | 28 | Angular_L |
| 32 | 4.59 | 39 | 13 | -16 | Insula_R |
| 36 | 4.58 | 13 | 30 | 32 | Cingulum_Mid_R |
| 37 | 4.57 | 6 | 40 | 19 | Cingulum_Ant_R |
| ***Girls*** |  |  |  |  |  |
| 493 | 5.52 | 17 | -82 | -1 | Lingual_R, Calcarine_R |
| 1931 | 5.29 | 39 | 8 | -47 | Temporal_Inf_R, Temporal_Pole_Mid |
| 4156 | 5.28 | -2 | -67 | 20 | Calcarine_L, Cuneus_L |
| 907 | 5.28 | 46 | 14 | -30 | Temporal_Pole_Mid_R |
| 1160 | 5.24 | -24 | -72 | -15 | Fusiform_L |
| 429 | 5.23 | -26 | -97 | 7 | Occipital_Mid_L |
| 154 | 5.15 | -33 | -69 | -8 | Fusiform_L |
| 341 | 5.12 | -53 | -64 | 19 | Temporal_Mid_L |
| 586 | 5.12 | -22 | -86 | 46 | Occipital_Sup_L |
| 918 | 5.08 | -32 | -1 | -44 | Temporal_Inf_L, Fusiform_L |
| 265 | 4.95 | 59 | -51 | 36 | Angular_R |
| 162 | 4.86 | -9 | -98 | -3 | Calcarine_L |
| 190 | 4.85 | 8 | -10 | 61 | Supp_Motor_Area_R |
| 269 | 4.85 | 9 | 15 | -3 | Caudate_R |
| 108 | 4.76 | -42 | 7 | -27 | Temporal_Mid_L |
| 83 | 4.74 | 69 | -30 | 27 | SupraMarginal_R |
| 242 | 4.73 | -17 | -31 | -19 | Cerebelum_4_5_L, ParaHippocampal_L |
| 151 | 4.72 | 2 | -94 | 16 | Cuneus_L |
| 39 | 4.7 | 62 | -54 | 24 | Temporal_Sup_R |
| 40 | 4.69 | 51 | 11 | 14 | Frontal_Inf_Oper_R |
| 74 | 4.66 | 22 | -54 | 12 | Calcarine_R |
| 39 | 4.63 | 59 | -56 | 12 | Temporal_Mid_R |
| 53 | 4.58 | -51 | 3 | -20 | Temporal_Mid_L |
| ***Boys*** |  |  |  |  |  |
| 8562 | 6.2 | 23 | 20 | -10 | Frontal_Inf_Orb_R, Caudate_R, Putamen_R |
| 2444 | 6.18 | -25 | -21 | -24 | ParaHippocampal_L |
| 5118 | 6.14 | 63 | 13 | 30 | Precentral_R |
| 2286 | 5.74 | -23 | 18 | 3 | Putamen_L, Insula_L |
| 2024 | 5.69 | 27 | -95 | 29 | Occipital_Inf_R |
| 1498 | 5.64 | -26 | 28 | 54 | Frontal_Mid_L |
| 691 | 5.58 | 28 | -58 | -14 | Fusiform_R, Lingual_R |
| 1371 | 5.57 | 25 | -11 | -23 | ParaHippocampal_R |
| 1393 | 5.51 | -20 | -65 | -10 | Lingual_L, Fusiform_L |
| 1283 | 5.51 | -4 | -66 | 10 | Lingual_L |
| 1869 | 5.36 | 6 | 63 | -4 | Frontal_Med_Orb, Frontal_Sup_Medial |
| 2020 | 5.32 | -22 | -76 | -53 | Cerebelum_8_L, Cerebelum_Crus2_L |
| 1865 | 5.28 | -37 | -24 | 39 | Postcentral_L |
| 730 | 5.21 | 20 | -80 | -50 | Cerebelum_7b_R, Cerebelum_Crus2_R |
| 667 | 5.17 | -27 | -63 | -30 | Cerebelum_6_L |
| 766 | 5.12 | 18 | -16 | 12 | Thalamus_R |
| 1001 | 5.09 | -38 | -90 | 20 | Occipital_Mid_L |
| 2177 | 5.06 | -67 | -26 | 17 | Temporal_Sup_L, Temporal_Mid_L |
| 171 | 4.93 | 38 | -72 | -47 | Cerebelum_Crus2_R |
| 58 | 4.86 | 23 | -37 | -10 | ParaHippocampal_R |
| 115 | 4.85 | -7 | -62 | 57 | Precuneus_L |
| 143 | 4.83 | -30 | -56 | -50 | Cerebelum_8_L |
| 125 | 4.83 | -9 | 56 | 21 | Frontal_Sup_Medial_L |
| 57 | 4.82 | 39 | -30 | -20 | Fusiform_R |
| 116 | 4.81 | 28 | -62 | -30 | Cerebelum_6_R |
| 146 | 4.78 | 31 | 64 | 6 | Frontal_Mid_R |
| 214 | 4.77 | -56 | 3 | -22 | Temporal_Mid_L |
| 169 | 4.75 | -20 | 44 | -17 | Frontal_Sup_Orb_L |
| 142 | 4.73 | 49 | 40 | 15 | Frontal_Inf_Tri_R |
| 117 | 4.72 | -34 | 39 | -14 | Frontal_Mid_Orb_L |
| 78 | 4.69 | 33 | 59 | 21 | Frontal_Mid_R |
| 88 | 4.67 | -30 | 58 | -2 | Frontal_Sup_Orb_L |
| 34 | 4.64 | 34 | 53 | 31 | Frontal_Mid_R |

*Note*: All clusters showed negative correlations; brain regions are identified with the Automated Anatomic Labeling atlas (Tzourio-Mazoyer et al., 2002). The sign of Z value indicates the direction of correlation. R: right; L: left; sup: superior; mid: middle; inf: inferior; G: gyrus

**Supplementary Table S6.** Slope tests of the correlation in ADHD scores between pairs of twins and unrelated children.

| **Variable** | **Pair** | **Slope-T** | **p-value** | **Correlations in MZ/DZ** |
| --- | --- | --- | --- | --- |
| ADHD score | MZ vs. DZ | 5.12 | 3.38e-07* | MZ: *r* = 0.61, *p* = 5.42e-34 |
| (all) | MZ vs. UR | 6.62 | 4.08e-11* | DZ: *r* = 0.23, *p* = 7.17e-19 |
|  | DZ vs. UR | 6.47 | 1.07e-10* |  |
| ADHD score | MZ vs. DZ | 2.26 | 2.39e-02* | MZ: *r* = 0.54, *p* = 2.77e-13 |
| (girls) | MZ vs. UR | 3.74 | 1.86e-04* | DZ: *r* = 0.27, *p* = 2.09e-09 |
|  | DZ vs. UR | 4.52 | 6.57e-06* |  |
| ADHD score | MZ vs. DZ | 4.92 | 1.08e-06* | MZ: *r* = 0.64, *p* = 1.16e-19 |
| (boys) | MZ vs. UR | 5.32 | 1.16e-07* | DZ: *r* = 0.23, *p* = 8.03e-08 |
|  | DZ vs. UR | 3.77 | 1.68e-04* |  |

*Note:* MZ: monozygotic; DZ: dizygotic; UR: unrelated. **p* < 0.05.

**Supplementary Table S7.** Slope tests of the correlation in GMV correlates between pairs of twins and unrelated children.

| **Variable** | **Pair** | **Slope-T** | **p-value** | **Correlations in MZ/DZ** |
| --- | --- | --- | --- | --- |
| ADHD – positive GMV correlate (HT) for all | MZ vs. DZ | 3.86 | 1.16e-04* | MZ: *r* = 0.72, *p* = 1.76e-51 |
|  | MZ vs. UR | 10.72 | 0.00e+00* | DZ: *r* = 0.46, *p* = 1.10e-74 |
|  | DZ vs. UR | 15.05 | 0.00e+00* |  |
| ADHD – negative GMV correlates for all | MZ vs. DZ | 9.10 | 0.00e+00* | MZ: *r* = 0.94, *p* = 1.62e-150 |
|  | MZ vs. UR | 17.32 | 0.00e+00* | DZ: *r* = 0.54, *p* = 4.28e-107 |
|  | DZ vs. UR | 18.06 | 0.00e+00* |  |
| ADHD – negative girls-specific GMV correlates | MZ vs. DZ | 6.72 | 4.02e-11* | MZ: *r* = 0.93, *p* = 2.97e-70 |
|  | MZ vs. UR | 12.93 | 0.00e+00* | DZ: *r* = 0.59, *p* = 5.94e-46 |
|  | DZ vs. UR | 12.00 | 0.00e+00* |  |
| ADHD – negative boys-specific GMV correlates | MZ vs. DZ | 5.07 | 5.14e-07* | MZ: *r* = 0.93, *p* = 1.13e-66 |
|  | MZ vs. UR | 11.34 | 0.00e+00* | DZ: *r* = 0.60, *p* = 4.64e-52 |
|  | DZ vs. UR | 12.74 | 0.00e+00* |  |

*Note:* MZ: monozygotic; DZ: dizygotic; UR: unrelated. **p* < 0.05. HT: hypothalamus

**Supplementary Table S8**. Univariate genetic model fitting of ADHD scores, GMV correlates, and 2-back%.

| *Variable* | *χ*^2^/*df* | RMSEA | TLI |
| --- | --- | --- | --- |
| ADHD score (all) | 14.79 | 0.13 | 0.87 |
| ADHD score (girls) | 6.76 | 0.19 | 0.77 |
| ADHD score (boys) | 12.41 | 0.19 | 0.73 |
| GMV positive correlate (all, HT) | 3.53 | 0.05 | 0.98 |
| GMV negative correlate (all) | 9.68 | 0.10 | 0.97 |
| GMV correlate (“girls specific”) | 7.39 | 0.09 | 0.98 |
| GMV correlate (“boys specific”) | 8.18 | 0.09 | 0.98 |
| 2-back% (all) | 1.94 | 0.04 | 0.98 |
| 2-back% (girls) | 0.33 | 0.00 | 1.00 |
| 2-back% (boys) | 3.31 | 0.10 | 0.89 |

*Note*: *df*: degrees of freedom; RMSEA: root-mean-square error of approximation; TLI: Tucker-Lewis Index. A *χ*^2^/df less than 2, RMSEA less than 0.06, or TLI greater than 0.95 indicate good model fit. Age, sex (for “all”), race, and study site were included as covariates for ADHD scores and 2-back%. Scanner model and TIV were included as additional covariates for GMV correlates.

**Supplementary Table S9a**. Univariate genetic model fitting of caudate, putamen and pallidum (AAL masks) GMV in girls and boys combined and separately.

| **Region** | ***χ*^2^/*df*** | **RMSEA** | **TLI** |
| --- | --- | --- | --- |
| ***All*** |  |  |  |
| caudate (L+R) | 1.83 | 0.03 | 1.00 |
| caudate_L | 1.37 | 0.02 | 1.00 |
| caudate_R | 2.10 | 0.04 | 1.00 |
| putamen (L+R) | 2.18 | 0.04 | 1.00 |
| putamen_L | 2.04 | 0.04 | 1.00 |
| putamen_R | 2.22 | 0.04 | 1.00 |
| pallidum (L+R) | 2.18 | 0.04 | 1.00 |
| pallidum_L | 1.73 | 0.03 | 1.00 |
| pallidum_R | 2.43 | 0.04 | 0.99 |
| ***Girls*** |  |  |  |
| caudate (L+R) | 1.71 | 0.05 | 1.00 |
| caudate_L | 1.05 | 0.01 | 1.00 |
| caudate_R | 2.00 | 0.06 | 0.99 |
| putamen (L+R) | 2.95 | 0.08 | 0.99 |
| putamen_L | 2.14 | 0.06 | 0.99 |
| putamen_R | 3.06 | 0.08 | 0.99 |
| pallidum (L+R) | 0.85 | 0.00 | 1.00 |
| pallidum_L | 0.95 | 0.00 | 1.00 |
| pallidum_R | 0.65 | 0.00 | 1.00 |
| ***Boys*** |  |  |  |
| caudate (L+R) | 1.29 | 0.05 | 1.00 |
| caudate_L | 1.52 | 0.03 | 1.00 |
| caudate_R | 1.08 | 0.02 | 1.00 |
| putamen (L+R) | 1.89 | 0.05 | 0.99 |
| putamen_L | 2.36 | 0.06 | 0.99 |
| putamen_R | 1.27 | 0.03 | 1.00 |
| pallidum (L+R) | 1.42 | 0.04 | 1.00 |
| pallidum_L | 1.63 | 0.04 | 0.99 |
| pallidum_R | 0.86 | 0.00 | 1.00 |

*Note*: *df*: degrees of freedom; RMSEA: root-mean-square error of approximation; TLI: Tucker-Lewis Index. A *χ*^2^/df less than 2, RMSEA less than 0.06, or TLI greater than 0.95 indicate good model fit. Age, sex (for “all”), race, and study site were included as covariates for ADHD scores and 2-back%. Scanner model and TIV were included as additional covariates for GMV correlates.

**Supplementary Table S9b.** Genetic, shared environmental, and non-shared environmental effects of caudate, putamen and pallidum (AAL masks) GMV in girls and boys combined and separately.

| **Region** | ***a*^2^ (A)** | ***c*^2^ (C)** | ***e*^2^ (E)** |
| --- | --- | --- | --- |
| ***All*** |  |  |  |
| caudate (L+R) | 0.90 [0.88, 0.92] | 0.00 [0.00, 0.00] | 0.10 [0.09, 0.12] |
| caudate_L | 0.88 [0.86, 0.90] | 0.00 [0.00, 0.00] | 0.12 [0.10, 0.14] |
| caudate_R | 0.88 [0.80, 0.90] | 0.00 [-0.08, 0.08] | 0.12[0.10, 0.14] |
| putamen (L+R) | 0.90 [0.88, 0.91] | 0.00 [0.00, 0.00] | 0.10 [0.09, 0.12] |
| putamen_L | 0.88 [0.86, 0.90] | 0.00 [0.00, 0.00] | 0.12 [0.10, 0.14] |
| putamen_R | 0.87 [0.85, 0.89] | 0.00 [0.00, 0.00] | 0.13 [0.11, 0.15] |
| pallidum (L+R) | 0.83 [0.80, 0.86] | 0.00 [0.00, 0.00] | 0.17 [0.14, 0.20] |
| pallidum_L | 0.79 [0.76, 0.83] | 0.00 [0.00, 0.00] | 0.21 [0.17, 0.24] |
| pallidum_R | 0.80 [0.77, 0.84] | 0.00 [0.00, 0.00] | 0.20 [0.17, 0.23] |
| ***Girls*** |  |  |  |
| caudate (L+R) | 0.84 [0.70, 0.99] | 0.03 [-0.11, 0.17] | 0.13 [0.10, 0.15] |
| caudate_L | 0.84 [0.69, 0.99] | 0.02 [-0.13, 0.16] | 0.14 [0.11, 0.18] |
| caudate_R | 0.82 [0.67, 0.96] | 0.05 [-0.09, 0.19] | 0.13 [0.11, 0.17] |
| putamen (L+R) | 0.87 [0.84, 0.90] | 0.00 [0.00, 0.00] | 0.13 [0.10, 0.16] |
| putamen_L | 0.85 [0.82, 0.89] | 0.00 [0.00, 0.00] | 0.15 [0.11, 0.18] |
| putamen_R | 0.85 [0.82, 0.89] | 0.00 [0.00, 0.00] | 0.15 [0.12, 0.18] |
| pallidum (L+R) | 0.83 [0.79, 0.87] | 0.00 [0.00, 0.00] | 0.17 [0.13, 0.21] |
| pallidum_L | 0.79 [0.74, 0.84] | 0.00 [0.00, 0.00] | 0.21 [0.16, 0.26] |
| pallidum_R | 0.81 [0.77, 0.86] | 0.00 [0.00, 0.00] | 0.19 [0.14, 0.23] |
| ***Boys*** |  |  |  |
| caudate (L+R) | 0.83 [0.70, 0.97] | 0.07 [-0.06, 0.20] | 0.10 [0.08, 0.12] |
| caudate_L | 0.82 [0.68, 0.96] | 0.06 [-0.08, 0.30] | 0.12 [0.10, 0.15] |
| caudate_R | 0.79 [0.66, 0.92] | 0.09 [-0.04, 0.22] | 0.12 [0.09, 0.15] |
| putamen (L+R) | 0.89 [0.86, 0.92] | 0.00 [0.00, 0.00] | 0.11 [0.08, 0.14] |
| putamen_L | 0.87 [0.84, 0.90] | 0.00 [0.00, 0.00] | 0.13 [0.10, 0.16] |
| putamen_R | 0.86 [0.83, 0.90] | 0.00 [0.00, 0.00] | 0.14 [0.10, 0.17] |
| pallidum (L+R) | 0.82 [0.78, 0.87] | 0.00 [0.00, 0.00] | 0.18 [0.14, 0.22] |
| pallidum_L | 0.78 [0.673, 0.83] | 0.00 [0.00, 0.00] | 0.22 [0.17, 0.27] |
| pallidum_R | 0.79 [0.74, 0.84] | 0.00 [0.00, 0.00] | 0.21 [0.16, 0.26] |

*Note*: L: left; R: right; *a*^2^: proportion of variance due to additive genetic effects (A); *c*^2^: proportion of variance due to shared environmental effects (C); *e*^2^: proportion of variance due to non-shared environmental effects (E); 95% confidence intervals are presented in the square brackets. Age, sex (for “all”), race, and study site were included as covariates for ADHD scores and 2-back%. Scanner model and TIV were included as additional covariates for GMV correlates.

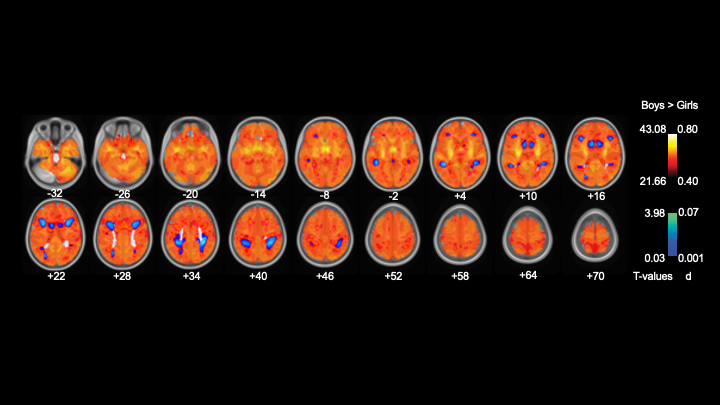

**Supplementary Figure S5a**. Unthresholded map of sex differences in GMVs. Color bars show the voxel *T* and the corresponding effect size (*d*) values.

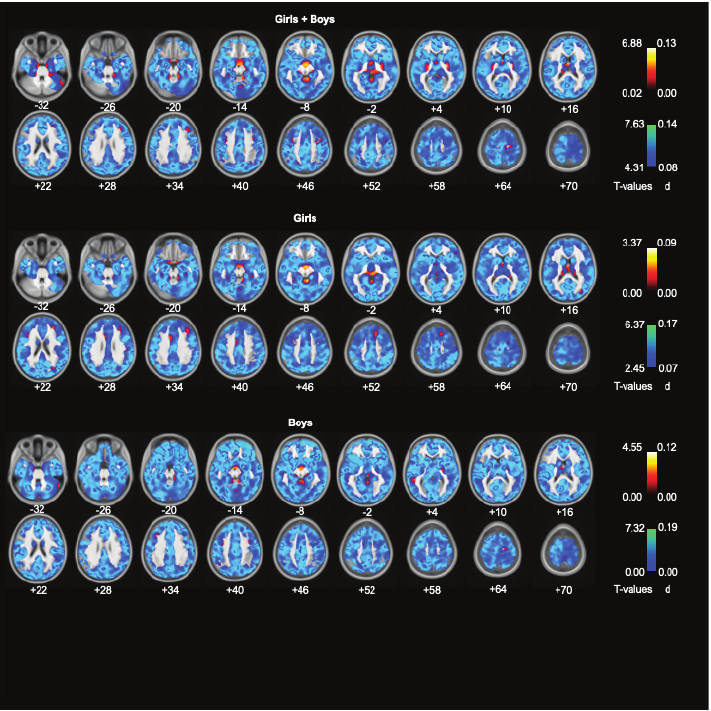

**Supplementary Figure S5b**. Unthresholded map of GMV correlates of ADHD score for girls and boys combined (top panels), and for girls (middle) and boys (bottom), separately. Color bars show voxel *T* values (left, cool colors to indicate negative correlation) and the corresponding effect sizes (right).

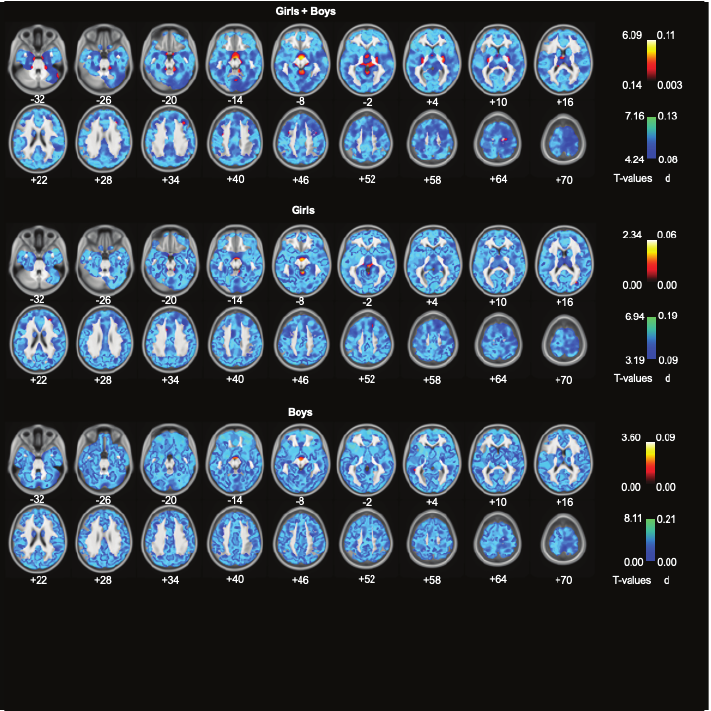

**Supplementary Figure S5c**. Unthresholded map of GMV correlates of ADHD scores for girls and boys combined, and for girls and boys, separately, without TIV included as a covariate. Cool color bars show voxel *T* (negative correlations) and *d* values.

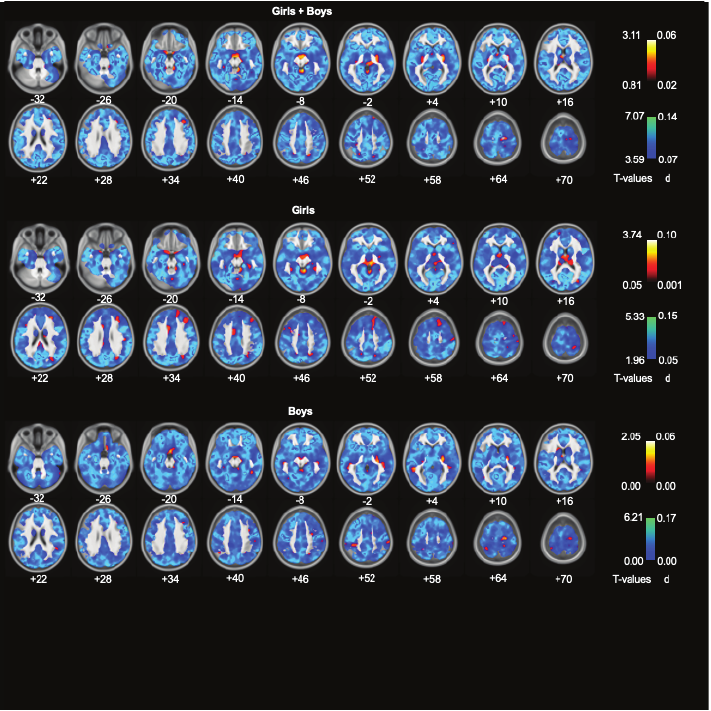

**Supplementary Figure S5d**. Unthresholded map of GMV correlates of ADHD scores (non-medicated children only) for girls and boys combined, and for girls and boys, separately. Cool color bars show voxel *T* (negative correlations) and *d* values.

**References**

Duvernoy, H.M., 2003. The human brain, 2nd ed. Springer-Verlag, Wien, Austria.

Tzourio-Mazoyer, N., Landeau, B., Papathanassiou, D., Crivello, F., Etard, O., Delcroix, N., Mazoyer, B., Joliot, M., 2002. Automated Anatomical Labeling of Activations in SPM Using a Macroscopic Anatomical Parcellation of the MNI MRI Single-Subject Brain. Neuroimage 15, 273-289.
